# Engineering eudicot-like lignin in rice via targeted disruption of grass-specific lignin modification pathways

**DOI:** 10.64898/2026.08.02.742291

**Authors:** Lydia Pui Ying Lam, Senri Yamamoto, Toshiaki Umezawa, Yuki Tobimatsu

## Abstract

- Lignin composition exhibits substantial diversity across plant lineages. Unlike eudicot and gymnosperm lignins, grass lignin incorporates *p*-coumarate and tricin units alongside canonical monolignols. To elucidate their specific biochemical and physiological roles, we engineered rice (*Oryza sativa*) to produce a “eudicot-like” lignin devoid of both modifications.
- Using CRISPR-Cas9, we generated mutants deficient in *p*-coumarate and tricin by simultaneously targeting their respective biosynthetic genes. The resulting mutant cell walls underwent structural and functional characterization via wet-chemical analyses, nuclear magnetic resonance, gel permeation chromatography, and antioxidant capacity assays.
- The newly generated *ospmt1/2 osfnsII* and *ospmt1/2 osa3′h/c5′h* triple-knockout mutants reached maturity with minor growth penalties. Subsequent cell wall analyses demonstrated near-complete depletion of both *p*-coumarate and tricin units in lignins. This structural shift substantially altered overall lignin content, monomeric composition, linkage distributions, and molecular weight, highlighting the divergent and synergistic roles of these units in lignin assembly and polymerization. Functionally, the free-radical scavenging capacity of rice lignin is markedly enhanced by *p*-coumaroylation, but attenuated by tricin incorporation.
- The successful synthesis of eudicot-like lignin within a grass system underscores the inherent plasticity of lignification. These engineered rice lines offer a valuable platform to investigate the physiological functions and biotechnological potential of grass lignin modifications.

## Introduction

Lignin is a complex, heterogeneous phenolic polymer deposited alongside cellulose and hemicellulose in the secondary cell walls of vascular plants, comprising approximately 10–30 wt% of typical plant biomass. It confers essential mechanical and biochemical properties to the cell wall, providing structural support, water impermeability for vascular transport, and resistance against UV and pathogen attack (Boerjan *et al*., 2003; Barros *et al*., 2015; Pesquet *et al*., 2025). While predominantly abundant in vascular tissues, lignification, i.e., deposition of lignin, occurs during the development of numerous specialized cell types, including endodermal and exodermal cells (particularly at Casparian strips), trichomes, abscission zones, and various reproductive organs (e.g., pollen, anthers, seed coats, and siliques) (Barros *et al*., 2015; Lee *et al*., 2018; Hao *et al*., 2024). Furthermore, ectopic lignification is rapidly triggered in response to a wide array of biotic and abiotic stresses (Barros *et al*., 2015; Cesarino, 2019; Lee *et al*., 2019). Beyond its biological functions, lignin is of paramount importance in modern biorefineries. As the most abundant aromatic biopolymer on Earth, it represents a promising renewable feedstock for synthesizing high-value aromatic chemicals and advanced materials (Ragauskas *et al*., 2014; Rinaldi *et al*., 2016; Schutyser *et al*. 2019; Abu-Omar *et al*., 2021; Arts *et al*., 2023). Paradoxically, however, lignin has long been recognized as the primary driver of biomass recalcitrance, impeding chemical pulping, forage digestibility, and the enzymatic release of fermentable sugars from lignocellulose (Ragauskas *et al*., 2014; Rinaldi *et al*., 2016). Alongside woody biomass, grass lignocellulose represents an abundant, sustainable resource for producing biochemicals and biofuels through biorefining (Davis *et al*., 2013; Tye *et al*., 2016; Bhatia *et al*., 2017; Umezawa *et al*., 2020). Yet, as discussed below, the structural complexity and structural diversity of grass cell wall components—particularly lignin—present distinct challenges for processing. This underscores the necessity of elucidating the biosynthesis and unique structural characteristics of grass lignin to optimize its industrial utility through targeted breeding and metabolic bioengineering (Halpin, 2019; Coomey *et al*., 2020; Umezawa *et al*., 2020; Chandrakanth *et al*., 2023; Peracchi *et al*., 2024; Umezawa, 2024).

Lignification in plant cell walls exhibits remarkable diversity regarding monomer assembly, yielding lignin polymers with compositions that are distinctive across plant lineages. In typical gymnosperms and eudicots—represented by softwoods and hardwoods, respectively—lignin is synthesized primarily through the oxidative radical coupling of three canonical monolignols: sinapyl, coniferyl, and *p*-coumaryl alcohols (Freudenberg, 1965; Sarkanen & Ludwig, 1971; Higuchi *et al*., 1985; Boerjan *et al*., 2003; Ralph *et al*., 2004, 2019). Upon polymerization, these precursors respectively form syringyl (S), guaiacyl (G), and *p*-hydroxyphenyl (H) lignin units (**Fig. 1a**). While gymnosperm lignin is composed primarily of G units, eudicot lignin comprises both G and S units; both polymer types contain minor amounts of H units. In contrast, the monomeric pool for monocot grass lignin is substantially more diverse. In addition to the canonical monolignols found in gymnosperms and eudicots, grass lignin incorporates monolignol *p*-coumarate conjugates (primarily sinapyl and coniferyl *p*-coumarates) (Ralph *et al*., 1994; Lu & Ralph, 1999; Hatfield *et al*., 2009; Ralph, 2010), and flavonoid tricin (del Río *et al*., 2012, 2020; Lan *et al*., 2015, 2016, 2025), producing the grass-specific tricin and *p*-coumarate modification units integrated to the lignin polymer (**Fig. 1a**). Minor amounts of monolignol ferulate conjugates are also incorporated, forming alkali-labile “zip-lignin” structures (Wilkerson *et al*., 2014; Karlen *et al*., 2016). Currently, the evolutionary, physiological, and biochemical significance of these grass-specific lignin modifications, and their impact on grass biomass utilization, remain poorly understood.

**Fig. 1.**
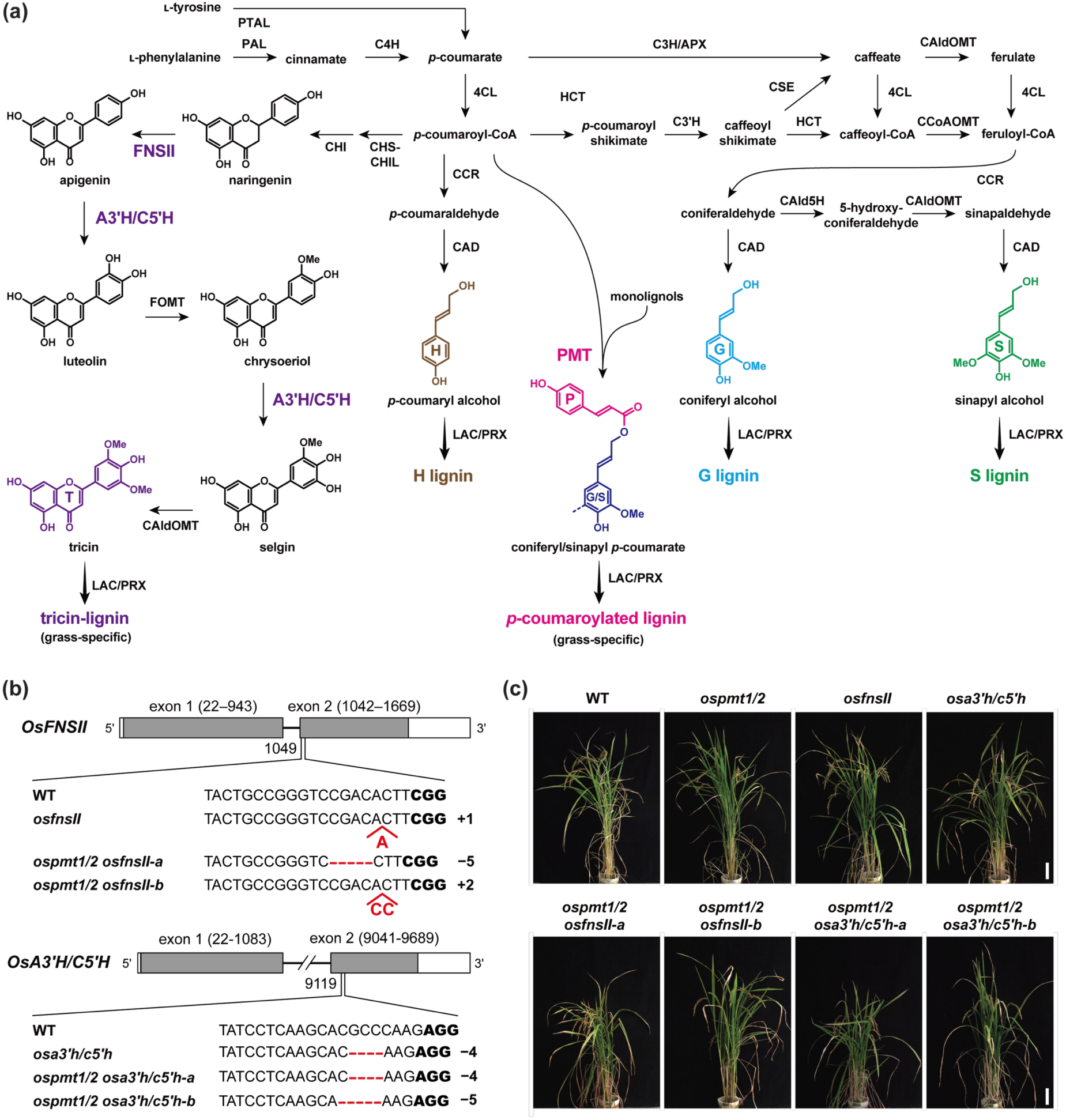
Grass lignin biosynthetic pathway and rice lignin mutants generated in this study. (a) Proposed lignin biosynthetic pathway in grasses. Grass lignin-specific enzymes targeted in this study, i.e., PMT, FNS and A3′H/C5′H, are highlighted. PTAL, ʟ-phenylalanine/ʟ-tyrosine ammonia-lyase; PAL, ʟ-phenylalanine ammonia-lyase; C4H, cinnamate 4-hydroxylase; 4CL, 4-hydroxycinnamate:CoA ligase; CAldOMT, 5-hydroxyconiferaldehyde *O*-methyltransferase; C3H, *p*-coumarate 3-hydroxylase; APX, ascorbate peroxidase; HCT, *p*-hydroxycinnamoyl-CoA:quinate/shikimate transferase; C3′H, *p*-coumaroyl ester 3-hydroxylase; CSE, caffeoyl shikimate esterase; CCoAOMT, caffeoyl-CoA *O*-methyltransferase; CAld5H, coniferaldehyde 5-hydroxylase; CCR, cinnamoyl-CoA reductase; CAD, cinnamyl alcohol dehydrogenase; PMT, *p*-coumaroyl-CoA:monolignol transferase; CHS, chalcone synthase; CHI, chalcone isomerase; CHIL, chalcone isomerase-like; FNSII, flavone synthase II; A3′H/C5′H, apigenin 3′-hydroxylase/chrysoeriol 5′-hydroxylase; F3ʹH, flavonoid 3ʹ-hydroxylase; FOMT, flavonoid *O*-methyltransferase; LAC, laccase; PRX, peroxidase. (b and c) Mutation patterns (b) and the mature-stage growth phenotypes (c) of the *p*-coumarate- and tricin-deficient genome-edited rice mutants generated in this study. In (b), deletions and insertions are highlighted in red, and the protospacer adjacent motif (PAM) sites are highlighted in bold. Mutation patterns in *OsPMT1* and *OsPMT2* are listed in Fig. S1. In (c), the scale bars indicate 10 cm. WT, wild type; *ospmt1/2*, *PMT*-deficient double-knockout line; *osfnsII*, *FNSII*-deficient single-knockout line; *osa3’h/c5’h*, *A3’H/C5’H*-deficient single-knockout line; *ospmt1/2 osfnsII-a/b*, *PMT*- and *FNSII*-deficient triple-knockout lines; *ospmt1/2 osa3’h/c5’h-a/b*, *PMT*- and *A3’H/C5’H*-deficient triple-knockout lines.

The biosynthetic pathways and polymerization mechanisms underlying the incorporation of *p*-coumarate and tricin units in grass lignin have been extensively investigated (Lam *et al*., 2021, 2023; Chandrakanth *et al*., 2023; Lan *et al*., 2025). The *p*-coumaroylation of monolignols is catalyzed by *p*-COUMAROYL-CoA:MONOLIGNOL TRANSFERASE (PMT), which facilitates the esterification of monolignols with *p*-coumaroyl-CoA (**Fig. 1a**) (Withers *et al*., 2012; Marita *et al*., 2014; Petrik *et al*., 2014; Lam *et al*., 2024b; Ji *et al*., 2025; Oliveira *et al*., 2025). The resulting monolignol *p*-coumarate conjugates are subsequently co-polymerized with non-acylated monolignols and tricin (Ralph *et al*., 1994; 2010; Lu & Ralph, 1999; Hatfield *et al*., 2008; Yamashita *et al*., 2020). Because *p*-coumarate moieties themselves do not readily participate in oxidative radical coupling during lignification, they remain as free-phenolic pendant groups decorating the lignin polymer (Hatfield *et al*., 2008; Ralph, 2010; Ralph *et al*., 2026).

In the tricin biosynthetic pathway, CHALCONE SYNTHASE (CHS) and CHALCONE ISOMERASE (CHI) divert metabolic flux from the phenylpropanoid pathway to produce the flavanone naringenin, the first intermediate possessing the canonical C_6_–C_3_–C_6_ flavonoid skeleton (**Fig. 1a**) (Eloy *et al*., 2017; Wang *et al*., 2020; Tetreault *et al*., 2021; Lam *et al*., 2022). The non-catalytic enhancer protein CHI-LIKE (CHIL) maximizes CHS activity (Ban *et al*., 2018; Waki *et al*., 2020; Lam *et al*., 2022; Imaizumi *et al*. 2026). Subsequently, FLAVONE SYNTHASE II (FNSII) desaturates naringenin to apigenin (Lam *et al*., 2014, 2017), followed by successive 3′- and 5′-hydroxylations and *O*-methylations. These downstream modifications are catalyzed by APIGENIN 3′-HYDROXYLASE/CHRYSOERIOL 5′-HYDROXYLASE (A3′H/C5′H) (Lam *et al*., 2015, 2019a) and either 5-HYDROXYCONIFERALDEHYDE *O*-METHYLTRANSFERASE (CAldOMT or COMT) or FLAVONOID OMT (FOMT) (**Fig. 1a**) (Kim *et al*., 2006; Lin *et al*., 2006; Lam et al., 2015, 2019b, 2024a; Fornalé *et al*., 2016; Eudes *et al*., 2017). Once biosynthesized, tricin is exported to the cell wall and incorporated into the lignin polymer via oxidative radical coupling, analogous to the polymerization of canonical monolignols (Lan *et al*., 2015, 2016a). Tricin cross-couples exclusively with monolignols or their *p*-coumarate conjugates and does not participate in subsequent chain propagation by coupling with the growing lignin polymer. As a result, tricin exclusively occupies the starting end of the final lignin polymer chain, functioning as a starter unit or a nucleation site in grass lignification (Lan *et al*., 2015; Berstis *et al*., 2021; Ralph *et al*., 2026).

Targeted disruption of the *p*-coumarate or tricin biosynthetic pathways has provided valuable insights into grass lignification, although their comprehensive structure–function relationships remain complex. In rice, maize (*Zea mays*), and *Brachypodium* (*Brachypodium distachyon*), disrupting *PMT* orthologs effectively eradicates *p*-coumarate decorations while frequently reducing the monolignol-derived S/G unit ratio (Marita *et al*., 2014; Petrik *et al*., 2014; Lam *et al*., 2024b; Ji *et al*., 2025; Oliveira *et al*., 2025). Conversely, silencing tricin biosynthetic genes (*CHS*, *CHI*, *CHIL*, *FNSII*, *A3ʹH/C5ʹH*, and *CAldOMT*) in rice, maize, and sorghum (*Sorghum bicolor*) successfully depletes lignin-bound tricin, occasionally causing the unnatural incorporation of intermediate flavonoids (e.g., naringenin and apigenin) or a concomitant reduction in S units primarily due to the bifunctional role of CAldOMT in tricin and S-type monolignol biosynthesis (Fornalé *et al*., 2016; Eloy *et al*., 2017; Eudes *et al*., 2017; Lam et al., 2017, 2019a, 2019b, 2022; Tetreault *et al*., 2021).

Despite these individual characterizations, the physiological and biochemical consequences of simultaneously eliminating both grass-specific lignin modification pathways remain unexplored. In this study, we engineered rice to deposit “eudicot-like” lignin completely devoid of both *p*-coumarate and tricin units by generating CRISPR-Cas9 multiplex knockout mutants simultaneously deficient in both lignin *p*-coumaroylation (*OsPMT1* and *OsPMT2*) and tricin biosynthesis (*OsFNSII* or *OsA3ʹH/C5ʹH*). We comprehensively evaluated the phenotypes of these stacked mutants alongside their corresponding parental lines deficient in either pathway, followed by in-depth structural characterization of their cell walls and isolated lignins using wet-chemical analyses, two-dimensional (2D) NMR spectroscopy, and gel permeation chromatography (GPC). Additionally, we functionally assessed their antioxidant capacities utilizing 2,2-diphenyl-1-picrylhydrazyl (DPPH) free-radical scavenging assays.

## Materials and Methods

### Plant materials

The *ospmt1/2* rice mutant was previously generated through CRISPR-Cas9-mediated knockout of *OsPMT1* and *OsPMT2* in the cv. Nipponbare background and is identical to the *ospmt1 ospmt2-2* line described by Lam et al. (2024b) (**Fig. S1**). To generate *FNSII*- and *A3ʹH/C5ʹH*-knockout mutants, single guide RNAs (sgRNAs) targeting the second exons of *OsFNSII* and *OsA3ʹH/C5ʹH* were designed using the CRISPR-P 2.0 program (Liu *et al*., 2017) (**Fig. 1b**) and integrated into the multiplex CRISPR-Cas9 binary vector pMgPoef4_129-2A-GFP (Toda *et al*., 2019; Yamamoto *et al*., 2024) using a Golden Gate Assembly Mix (New England Biolabs, USA) and the oligonucleotides listed in **Table S1**. The binary vectors were transformed into embryonic calli from wild-type rice (cv. Nipponbare) to generate *osfnsII* and *osa3ʹh/c5ʹh* single-knockout mutants, or into calli from *ospmt1/2* to generate *ospmt1/2 osfnsII* and *ospmt1/2 osa3ʹh/c5ʹh* triple-knockout mutants using the *Agrobacterium* strain EHA101 (Hiei *et al*., 1994). The regenerated T_0_ plants were genotyped and grown to maturity in potting soil in a growth chamber (Lam *et al*., 2017). Selected T_1_ plants were further genotyped and grown to maturity in a greenhouse maintained at 27 °C (Lam *et al*., 2017) along with wild-type (WT) rice and the *ospmt1/2* mutant line. Mature plants were phenotypically characterized (**Table 1**), harvested, and dried at room temperature for one week and at 45 °C for 3 days before further analyses. For genotyping, genomic DNA was extracted from young leaves, and the genomic region containing the target site was amplified by PCR using the primers listed in **Table S1**. The top three potential off-target sites predicted by CRISPR-P 2.0 (Liu *et al*., 2017) were analyzed by direct sequencing (**Table S2**) (Takeda et al., 2019a).

**Table 1.** Growth phenotype and biomass productivity of *p*-coumarate- and tricin-deficient rice mutants.

| traits | WT | *ospmt1/2* | *osfnsII* | *osa3'h/c5'h* | *ospmt1/2*<br>*osfnsII-a* | *ospmt1/2*<br>*osfnsII-b* | *ospmt1/2*<br>*osa3'h/c5'h-a* | *ospmt1/2*<br>*osa3'h/c5'h-b* |
| --- | --- | --- | --- | --- | --- | --- | --- | --- |
| plant height (cm)<sup>1</sup> | 114.3 ± 9.3<sup>a</sup> | 106.1 ± 5.2<sup>ab</sup> | 109.4 ± 3.7<sup>ab</sup> | 108.3 ± 3.2<sup>ab</sup> | **100.9 ± 6.2<sup>b</sup>** | 109.0 ± 3.2<sup>ab</sup> | **100.1 ± 7.8<sup>b</sup>** | **103.0 ± 3.0<sup>b</sup>** |
| culm length (cm)<sup>2</sup> | 82.8 ± 5.2<sup>a</sup> | **69.1 ± 2.9<sup>b</sup>** | **70.6 ± 3.9<sup>b</sup>** | **67.3 ± 6.0<sup>b</sup>** | **62.9 ± 3.7<sup>b</sup>** | **66.7 ± 3.6<sup>b</sup>** | **63.2 ± 5.7<sup>b</sup>** | **65.1 ± 0.8<sup>b</sup>** |
| panicle length (cm) | 19.8 ± 1.5<sup>ab</sup> | 17.7 ± 1.0<sup>ab</sup> | 19.4 ± 1.9<sup>ab</sup> | 20.8 ± 2.5<sup>a</sup> | **16.3 ± 2.5<sup>c</sup>** | 18.5 ± 1.7<sup>ab</sup> | **17.2 ± 1.1<sup>bc</sup>** | 17.9 ± 0.8<sup>ab</sup> |
| tiller number | 9.7 ± 2.2<sup>a</sup> | 8.8 ± 1.6<sup>a</sup> | 8.7 ± 2.5<sup>a</sup> | 8.3 ± 1.4<sup>a</sup> | 11.0 ± 2.8<sup>a</sup> | 9.5 ± 1.8<sup>a</sup> | 9.5 ± 2.3<sup>a</sup> | 9.0 ± 1.7<sup>a</sup> |
| panicle number | 9.8 ± 2.3<sup>a</sup> | 9.0 ± 1.5<sup>a</sup> | 9.0 ± 2.4<sup>a</sup> | 8.5 ± 1.5<sup>a</sup> | 11.2 ± 2.8<sup>a</sup> | 9.7 ± 2.1<sup>a</sup> | 10.5 ± 2.4<sup>a</sup> | 9.0 ± 1.7<sup>a</sup> |
| fertility rate (%) | 92.1 ± 4.5<sup>a</sup> | 87.2 ± 7.1<sup>ab</sup> | 86.7 ± 3.7<sup>ab</sup> | 89.1 ± 3.7<sup>ab</sup> | 81.0 ± 13.0<sup>ab</sup> | **78.3 ± 7.7<sup>b</sup>** | 84.3 ± 4.8<sup>ab</sup> | 82.5 ± 2.5<sup>ab</sup> |
| biomass (g)<sup>3</sup> | 29.7 ± 4.9<sup>a</sup> | **22.2 ± 3.5<sup>b</sup>** | **21.1 ± 3.5<sup>b</sup>** | **22.4 ± 3.5<sup>b</sup>** | **19.7 ± 3.6<sup>b</sup>** | **19.2 ± 2.4<sup>b</sup>** | **19.3 ± 2.6<sup>b</sup>** | **18.9 ± 3.6<sup>b</sup>** |

<sup>1</sup>Height between the base of the aerial part and the tip of the top leaf. <sup>2</sup>Height between the base of the aerial part to the base of panicle. <sup>3</sup>Dry weight of the whole aerial part excluding panicles. The values are means ± standard deviation from individually analyzed plants (*n* = 5). Different letters indicate significant differences (one-way ANOVA with Tukey HSD test, *P* < 0.05). Values highlighted in bold indicate significant differences from WT. WT, wild type; *ospmt1/2*, *PMT*-deficient double-knockout line; *osfnsII*, *FNSII*-deficient single-knockout line; *osa3'h/c5'h*, *A3'H/C5'H*-deficient single-knockout line; *ospmt1/2 osfnsII-a/b*, *PMT*- and *FNSII*-deficient triple-knockout lines; *ospmt1/2 osa3'h/c5'h-a/b*, *PMT*- and *A3'H/C5'H*-deficient triple-knockout lines.

### Cell wall and lignin sample preparations

Extractive-free cell wall residue (CWR) samples were prepared from dried mature rice culms as described previously (Yamamura *et al*., 2011). The dioxane/water-soluble lignin (DWL) samples used for NMR and GPC were prepared according to previously described methods (Martin *et al*., 2023; Lam *et al*., 2024b; Ji *et al*., 2025). In brief, CWR (∼1.5 g pooled across three biological replicates) was pulverized in zirconia vessels using a Pulverisette 7 planetary ball mill (Fritsch, Idar-Oberstein, Germany; 600 rpm, 36 cycles of 10 min milling and 5 min rest), and subsequently subjected to three cycles of enzymatic hydrolysis using crude cellulase (CELLULYSIN®, Merck Millipore, Darmstadt, Germany; 7.5 mg in 0.1 M sodium acetate buffer, pH 5.0). The digested CWR samples were then extracted with dioxane/water (96:4, v/v), and isolated material was purified by reprecipitation in 0.01 M aqueous hydrochloric acid, followed by washing with distilled water.

### Chemical analysis

The Klason lignin assay (Hatfield *et al*., 1994), analytical thioacidolysis (Yamamura *et al*., 2012; Yue *et al*., 2012), quantification of cell wall-bound *p*-coumarate and ferulate by mild alkaline hydrolysis (Yamamura *et al*., 2011), and neutral sugar analysis by two-step trifluoroacetic acid/sulfuric acid-catalyzed hydrolysis (Lam *et al*., 2017) were conducted according to previously described methods.

### 2D NMR analysis

For the NMR analysis of whole rice culm cell walls, approximately 200 mg of CWR samples (pooled from three biologically independent plants) were ball-milled using a Pulverisette 7 planetary ball mill (Fritsch, Idar-Oberstein, Germany) in zirconia vessels containing zirconia ball bearings (600 rpm, 10 cycles of 10 min milling with 5 min rests) (Yamamoto *et al*., 2024; Ji *et al*., 2025). Sixty milligrams of the ball-milled CWRs were directly swelled in 600 μL of dimethylsulfoxide-*d*_6_/pyridine-*d*_5_ (4:1, v/v) and subjected to ^1^H–^13^C heteronuclear single-quantum coherence (HSQC) NMR analysis (Kim & Ralph, 2010; Mansfield *et al*., 2012). For the NMR analysis of DWL samples, aliquots (approximately 20 mg) were completely dissolved in 500 μL of dimethyl sulfoxide-*d*_6_/pyridine-*d*_5_ (4:1, v/v) and then subjected to 2D HSQC NMR analysis (Martin *et al*., 2023; Lam *et al*., 2024b; Ji *et al*., 2025). The HSQC NMR spectra were acquired using a Bruker Avance III 800 system (800 MHz; Bruker Biospin, Billerica, MA, USA) equipped with a cryogenically cooled 5-mm TCI gradient probe. Adiabatic HSQC experiments were conducted using the standard Bruker implementation (‘hsqcetgpsp.3’) and previously described parameters (Kim & Ralph, 2010; Mansfield *et al*., 2012). Data were processed and analyzed using TopSpin 5.0 software (Bruker Biospin) as previously described (Martin *et al*., 2023; Lam *et al*., 2024; Yamamoto *et al*., 2024; Ji *et al*., 2025). Peak assignment was based on comparison with NMR data in the literature (Kim & Ralph, 2010; Mansfield *et al*., 2012; Lan *et al*., 2018; Lam *et al*., 2019a, 2024b; Ralph *et al*., 2024; Yamamoto *et al*., 2024; Ji *et al*., 2025). The central dimethyl sulfoxide solvent peaks (*δ*_C_/*δ*_H_, 39.5/2.49 ppm) were used for calibrating chemical shifts.

### GPC

The GPC analysis of the DWL samples was conducted following the method described previously (Ji *et al*., 2025). Briefly, the lignin samples were dissolved in *N*,*N*-dimethylformamide containing 0.1 M lithium bromide (approximately 3 mg mL^−1^) and subjected to GPC analysis on a Shimadzu LC-20AD system (Shimadzu, Kyoto, Japan) equipped with an SPD-20A UV/VIS detector using the following conditions: columns, Tosoh TSKgel α-M and α-2500; eluent, *N*,*N*-dimethylformamide containing 0.1 M lithium bromide; flow rate, 0.5 mL min^−1^; column oven temperature, 40 °C; sample injection volume, 20 µL; and detection, UV absorption at 280 nm. Data acquisition and processing were performed using LCsolution software (Shimadzu). Molecular weight calibration was conducted using Agilent EasyVials polystyrene standards (Agilent Technologies, Santa Clara, CA, USA).

### DPPH assay

The DPPH assay was conducted according to the method of Serpen *et al*. (2007) with minor modifications. The DPPH reagent was prepared by dissolving 5 mg of DPPH in 200 mL of methanol (Lin *et al*., 2021). For the assay, 10 mg of CWRs suspended in 20 µL of methanol was mixed with 1.7 mL of the DPPH reagent and incubated at room temperature for 30 min. The reaction mixtures were vortexed at 0, 15, and 25 min post-initiation. Following centrifugation (10,000 × *g*, 2 min, 4 °C), the absorbance of the supernatant was measured at 517 nm using a Shimadzu UV-2600 UV–Vis spectrophotometer (Shimadzu, Kyoto, Japan). A calibration curve was constructed in parallel using catechin standards (0–0.4 mg) under identical conditions. Antioxidant activities were calculated from this curve (Sawamura *et al*., 2017) and are expressed as catechin equivalent antioxidant capacity (CEAC) per gram of CWR or per gram of Klason lignin.

### Statistical analysis

One-way ANOVA followed by Tukey HSD multiple comparison test was performed using the GraphPad Prism version 8 (GraphPad Software, San Diego, CA, USA).

### Accession numbers

Sequence data from this article can be found under accession numbers: LOC_Os01g18744, *OsPMT1* (Withers *et al*., 2012); LOC_Os05g04584, *OsPMT2* (Lam *et al*., 2024b); LOC_Os04g01140, *OsFNSII* (Lam *et al*., 2014); LOC_Os10g16974, *OsA3ʹH*/*C5ʹH* (Lam *et al*., 2015).

## RESULTS

### Generation and phenotyping of *p*-coumarate- and tricin-deficient rice mutants

To eliminate *p*-coumarate and tricin units from rice lignin, we knocked out genes responsible for lignin *p*-coumaroylation, tricin biosynthesis, or both. As previously described, the *ospmt1/2* double-knockout mutant, which harbors mutations in both *OsPMT1* and *OsPMT2* (**Fig. S1**), completely lacks lignin-bound *p*-coumarate (Lam *et al*., 2024b). Using this *p*-coumarate-deficient background, we introduced additional null mutations into the key tricin biosynthetic genes *OsFNSII* (Lam *et al*., 2014, 2017) or *OsA3ʹH/C5ʹH* (Lam *et al*., 2015, 2021). As controls, corresponding single-knockout mutants for *OsFNSII* or *OsA3ʹH/C5ʹH* were generated in the same wild-type cultivar. Primary transformants (T_0_ generation) containing target-gene indels were identified via genotyping. From the T_1_ generation, we isolated four independent triple-homozygous mutant lines carrying distinct alleles of *OsFNSII* or *OsA3ʹH/C5ʹH* in addition to the *OsPMT1* and *OsPMT2* mutations (hereafter referred to as *ospmt1/2 osfnsII-a*, *ospmt1/2 osfnsII-b*, *ospmt1/2 osa3ʹh/c5ʹh-a*, and *ospmt1/2 osa3ʹh/c5ʹh-b*) (**Fig. 1b**). Additionally, two homozygous single mutants harboring indels in *OsFNSII* or *OsA3ʹH/C5ʹH* were isolated (hereafter named *osfnsII* and *osa3ʹh/c5ʹh*) (**Fig. 1b**). All identified indels caused frameshift mutations, resulting in complete gene knockouts (**Fig. S2,S3**). Sequence analysis of the top three predicted off-target sites confirmed the absence of off-target mutations (**Table S2**).

To evaluate phenotypic impacts, fully genotyped homozygous mutants (T_1_ generation) were grown alongside WT and *ospmt1/2* background plants and characterized at maturity (**Fig. 1c**; **Table 1**). As previously reported (Lam *et al*., 2017; 2019; 2024b), mutants deficient in *PMTs* (*ospmt1/2*), *FNSII* (*osfnsII*), or *A3ʹH/C5ʹH* (*osa3ʹh/c5ʹh*) exhibited mild growth reductions compared with WT plants, particularly in culm length and biomass productivity. Similarly, the newly developed *ospmt1/2 osfnsII* and *ospmt1/2 osa3ʹh/c5ʹh* triple-knockout lines displayed comparable penalties in culm length and biomass productivity. Additionally, the *ospmt1/2 osfnsII-a* and *ospmt1/2 osa3ʹh/c5ʹh* mutants exhibited slightly decreased plant height and/or panicle length. Fertility remained largely unaffected, with the exception of a marginal decrease in the *ospmt1/2 osfnsII-b* line. Taken together, these results indicate that while disrupting lignin *p*-coumaroylation and/or tricin biosynthetic genes results in minor growth reductions, it crucially does not induce severe developmental defects, even when both pathways are simultaneously disrupted.

### The *ospmt1/2 osfnsII* and *ospmt1/2 osa3ʹh/c5ʹh* rice produce “eudicot-like” lignins lacking *p*-coumarate and tricin units

To investigate how stacked mutations in lignin *p*-coumaroylation and tricin biosynthetic genes affect cell wall architecture, extractive-free CWR samples from mature culms of mutant and WT plants were analyzed using chemical and NMR methods.

#### Chemical analyses

We first examined lignocellulose composition—specifically, the content and composition of lignin, polysaccharides, and cell wall-bound hydroxycinnamates (*p*-coumarate and ferulate). Klason analysis revealed no significant changes in lignin content between the WT and *ospmt1/2* plants. In contrast, all lines deficient in *FNSII* or *A3ʹH/C5ʹH* (*osfnsII*, and *osa3ʹh/c5ʹh osa3ʹh/c5ʹh*) displayed significantly reduced lignin content (reduced by ∼24–12% compared to WT) (**Fig. 2a**). No further reductions were observed in the triple mutants (*ospmt1/2 osfnsII* and *ospmt1/2 osa3ʹh/c5ʹh*) compared with their respective tricin single-knockout mutants (*osfnsII* and *osa3ʹh/c5ʹh*). Consistent with the Klason lignin data, the total yield of lignin monomers released by thioacidolysis was significantly reduced (by ∼22– 33% compared to WT) in all lines deficient in *FNSII* or *A3ʹH/C5ʹH* (*osfnsII*, *osa3ʹh/c5ʹh*, *ospmt1/2 osfnsII*, and *ospmt1/2 osa3ʹh/c5ʹh*), while increased (by ∼24%) in the *PMT*-deficient *ospmt1/2* line (**Fig. 2b**). These results demonstrate that disrupting tricin-associated *FNSII* and *A3ʹH/C5ʹH* genes, but not *p*-coumarate-associated *PMT* genes, significantly reduces total lignin content.

**Fig. 2.**
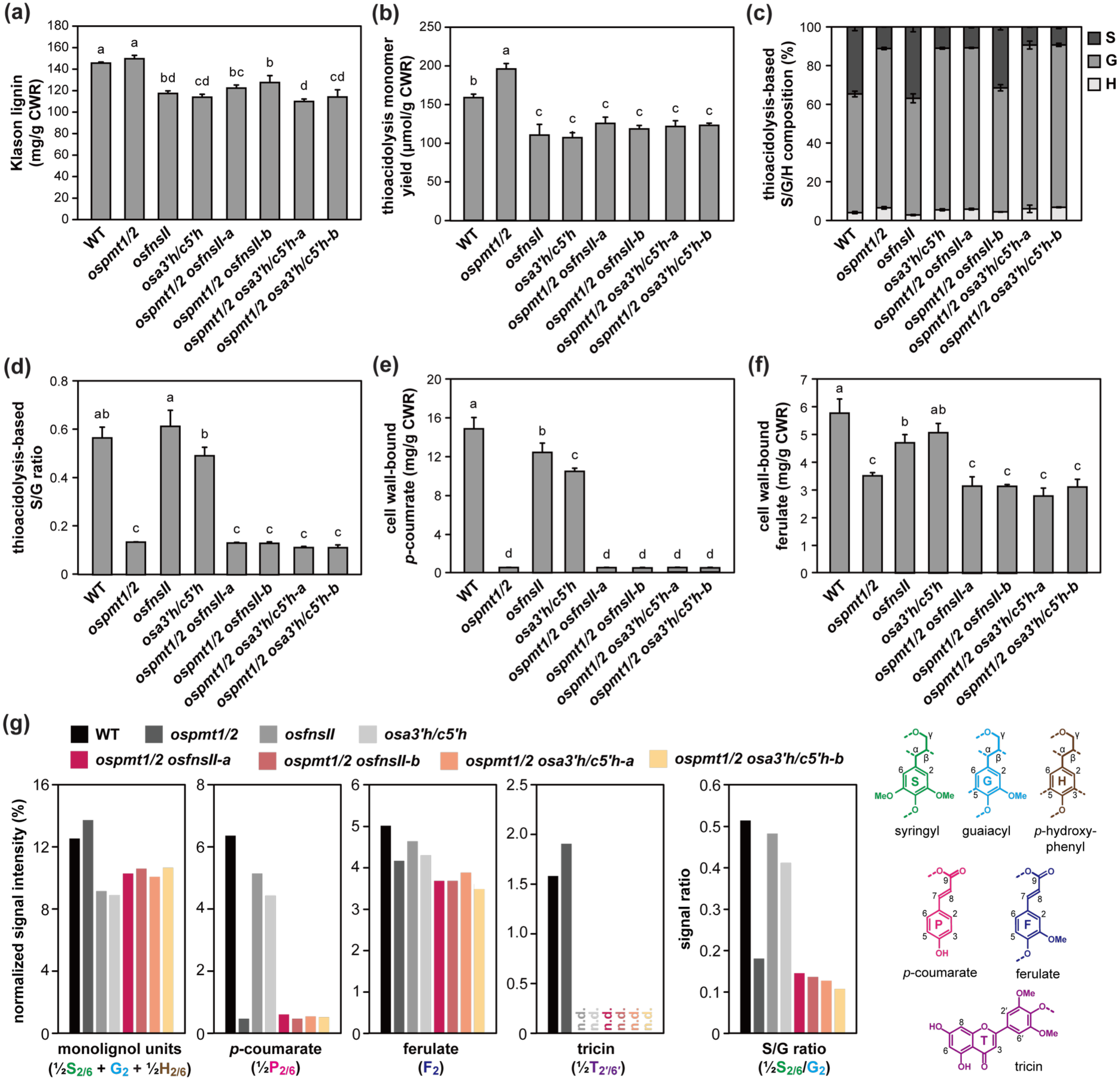
Lignin content and composition analyses of *p*-coumarate- and tricin-deficient rice mutant cell walls. (a-f) Chemical analysis data of culm cell wall residue (CWR) samples prepared from rice mutant lines. Klason lignin content (a), thioacidolysis-derived total lignin monomer yield (b), monomer composition (c) and S/G monomer ratio (d), and mild alkaline hydrolysis-derived cell-wall-bound *p*-coumarate (e) and ferulate (f) contents are reported. The values are means ± standard deviation from individually analyzed plants (*n* = 3). Different letters indicate significant differences (one-way ANOVA with Tukey HSD test, *P* < 0.05). (g) Normalized signal intensities of major aromatic unit signals in 2D HSQC NMR spectra of whole rice culm cell walls. NMR analysis was conducted for culm CWR samples pooled from three biologically independent plants for each line. The cell wall NMR spectra, complete intergradation data, and peak assignments are shown in **Figs. S4** and **S5**, and **Table S4**. n.d., not detected; WT, wild type; *ospmt1/2*, *PMT*-deficient double-knockout line; *osfnsII*, *FNSII*-deficient single-knockout line; *osa3’h/c5’h*, *A3’H/C5’H*-deficient single-knockout line; *ospmt1/2 osfnsII-a/b*, *PMT*- and *FNSII*-deficient triple-knockout lines; *ospmt1/2 osa3’h/c5’h-a/b*, *PMT*- and *A3’H/C5’H*-deficient triple-knockout lines.

Analytical thioacidolysis further revealed changes in monolignol unit composition across the mutants. Notably, all lines deficient in *PMTs* (*ospmt1/2*, *ospmt1/2 osfnsII*, and *ospmt1/2 osa3ʹh/c5ʹh*) displayed a marked reduction (by ∼69–74% compared to WT) in the proportion of S lignin units, whereas mutants deficient only in *FNSII* (*osfnsII*) or *A3ʹH/C5ʹH* (*osa3ʹh/c5ʹh*) retained an S/G/H unit composition comparable to that of WT rice (**Fig. 2c,d**). These results corroborate our earlier findings that the incorporation of S-type monolignols into grass lignin is closely associated with monolignol *p*-coumaroylation catalyzed by PMT (Lam *et al*., 2024b; Ji *et al*., 2025; Oliveira *et al*., 2025).

We quantified cell wall-bound *p*-coumarate and ferulate following their release via mild alkaline hydrolysis. Consistent with the established role of PMT in lignin *p*-coumaroylation, *p*-coumarate was drastically depleted in all *PMT*-deficient lines (*ospmt1/2*, *ospmt1/2 osfnsII*, and *ospmt1/2 osa3ʹh/c5ʹh*) (**Fig. 2e**). The trace amounts of *p*-coumarate detected in these mutants likely originate from the *p*-coumaroylation of hemicelluloses (mainly arabinoxylan) (Ralph, 2010; Lam *et al*., 2024b; Yamamoto *et al*., 2024), as confirmed by our subsequent analysis of isolated lignins. Furthermore, *p*-coumarate content was also significantly reduced (by 18–30% compared to WT) in the tricin single-knockout mutants (*osfnsII* and *osa3ʹh/c5ʹh*). Because *p*-coumarate is primarily esterified to S lignin in grass cell walls including in rice (Ralph, 2010; Lam *et al*., 2024b; Yamamoto *et al*., 2024), these reductions likely attributable to the overall reduction in lignin, particularly S units (**Fig. 2a-c**). Ferulate content was moderately reduced (by ∼38–48%) across all *p*-coumarate-deficient mutants (*ospmt1/2*, *ospmt1/2 osfnsII*, and *ospmt1/2 osa3ʹh/c5ʹh*), whereas it was only slightly reduced or unchanged in the tricin single-knockout lines (*osfnsII* and *osa3ʹh/c5ʹh*) (**Fig. 2f**). These results suggest that disrupting lignin *p*-coumaroylation may also affect cell wall feruloylation, as reported previously (Petrik *et al*., 2014; Lam *et al*., 2024b; Ji *et al*., 2025; Oliveira *et al*., 2025).

Neutral sugar analysis revealed slight increases in glucose released from crysltalline cellulose across all *p*-coumarate- and tricin-deficient mutants, with the greatest increases observed in the *ospmt1/2 osfnsII* and *ospmt1/2 osa3ʹh/c5ʹh* triple-knockout mutants (**Table S3**). Furthermore, galactose content was slightly elevated across all tricin-deficient lines (*osfnsII*, *osa3ʹh/c5ʹh*, *ospmt1/2 osfnsII*, and *ospmt1/2 osa3ʹh/c5ʹh*). The levels of all other neutral sugars remained largely comparable to those of WT plants, with the exception of minor increases in amorphous glucose in the *osfnsII* mutant and mannose in the *ospmt1/2 osfnsII-b ospmt1/2 osa3ʹh/c5ʹh-b* mutants.

#### Cell wall NMR analysis

To further investigate shifts in lignin aromatic composition across the mutants, we analyzed ball-milled culm CWRs using whole-cell-wall 2D HSQC NMR spectroscopy in a dimethylsulfoxide-*d*_6_/pyridine-*d*_5_ solvent system (Kim & Ralph, 2010; Mansfield *et al*., 2012). The resulting spectra displayed characteristic signals from major lignin aromatic units—including S, G, and H units (**S**, **G**, and **H**) consisting the monolignol-derived polymer backbone, as well as *p*-coumarate (**P**), ferulate (**F**), and tricin (**T**) units typical of grass lignins (**Fig. S4; Table S4**). For semi-quantitative comparison, we performed volume integration of well-resolved lignin and *p*-hydroxycinnamate aromatic signals (**½S_2/6_**, **G_2_**, **½H_2/6_**, **½P_2/6_**, **F_2_**, and **½T_2ʹ/6ʹ_**) along with the major polysaccharide anomeric signals (**Gl_1_**, **X_1_**, **Xʹ_1_**, **Xʹʹ_1_**, **A_1_**, **Ga_1_**, and **U_1_**) (**Fig. S4; Table S4**). Relative signal intensities were normalized to the sum of these integrated contours, approximately reflecting the proportional abundance of each component in the cell walls (Yamamoto *et al*., 2024; Ji *et al*., 2025) (**Fig. 2g; Fig. S5**).

The obtained volume integration data further corroborated the lignin compositional changes observed following disruption of the lignin *p*-coumaroylation and tricin pathways (**Fig. 2g**). Consistent with our Klason lignin and thioacidolysis data, the sum of the S/G/H unit signals (**½S_2/6_** + **G_2_** + ½**H_2/6_**) was reduced across all tricin-deficient lines (*osfnsII*, *osa3ʹh/c5ʹh*, *ospmt1/2 osfnsII*, and *ospmt1/2 osa3ʹh/c5ʹh*), indicating a concomitant decrease in total cell wall lignin upon the disruption of tricin biosynthesis. Furthermore, mirroring our thioacidolysis and mild alkaline hydrolysis results, the S/G signal ratio (**½S_2/6_**/**G_2_**) and the *p*-coumarate signal (**½P_2/6_**) were markedly lower across all *PMT*-deficient backgrounds (*ospmt1/2*, *ospmt1/2 osfnsII*, and *ospmt1/2 osa3ʹh/c5ʹh*). Finally, the tricin signal (**½T_2ʹ/6ʹ_**) was completely absent in all tricin-deficient lines, demonstrating the successful elimination of lignin-bound tricin. Trace novel flavonoid signals corresponding to naringenin incorporated into lignin (Lam *et al*., 2017) were also detected in the cell wall NMR spectra of the *FNSII*-deficient *osfnsII* and *ospmt1/2 osfnsII* mutants (**Fig. S4**); these will be explored in greater detail during our subsequent analysis of isolated lignins.

Collectively, our chemical and NMR data demonstrate that introducing stacked mutations in *p*-coumarate and tricin pathway genes resulted in a “eudicot-like” cell wall lignin in rice—one predominantly composed of canonical monolignol-derived S/G/H units and virtually devoid of grass-specific *p*-coumarate and tricin modifications.

### In-depth structural characterizations of eudicot-like lignins produced by *p*-coumarate- and tricin-deficient rice

To elucidate the structural features of eudicot-like lignins produced by the *ospmt1/2 osfnsII* and *ospmt1/2 osa3ʹh/c5ʹh* rice mutants, DWL (dioxane/water-soluble lignin) samples isolated from culm CWR were subjected to additional lignin structural analysis using 2D HSQC NMR and GPC. Utilizing these polysaccharide-depleted, soluble DWL samples facilitated a rigorous assessment of lignin linkage and molecular weight distributions—details that remain largely inaccessible via the chemical and 2D NMR analyses of intact cell wall samples described above. For comparison, DWL samples prepared from the WT control, *ospmt1/2*, *osfnsII* and *osa3ʹh/c5ʹh* lines were evaluated in parallel.

#### Aromatic composition

Analysis of the aromatic sub-regions (*δ*_C_/*δ*_H_, 150–90/8.0–6.0 ppm) of the 2D HSQC NMR spectra further unequivocally established the eudicot-like aromatic composition of the *ospmt1/2 osfnsII* and *ospmt1/2 osa3ʹh/c5ʹh* rice mutant lignins (**Fig. 3a; Table S5**). To evaluate variations in aromatic unit composition across the isolated lignins, volume integrals were normalized as percentages relative to the sum of the major S/G/H aromatic signals (**½S**_2/6_ + **G**_2_ + **½H**_2/6_ = 100%) (**Fig. 3b,c**). In agreement with their genetic backgrounds, *p*-coumarate signals (**P**) were completely absent in all *PMT*-deficient lines (*ospmt1/2*, *ospmt1/2 osfnsII*, and *ospmt1/2 osa3ʹh/c5ʹh*), whereas tricin signals (**T**) were undetectable across all *FNSII-* and *A3ʹH/C5ʹH*-deficient lines (*osfnsII*, *osa3ʹh/c5ʹh*, *ospmt1/2 osfnsII*, and *ospmt1/2 osa3ʹh/c5ʹh*) (**Fig. 3a,c**). The absence of *p*-coumarate in isolated lignins from *PMT*-deficient lines confirmed that the trace *p*-coumarate detected in their cell walls (**Fig. 2**) is associated with hemicelluloses rather than lignin. Consistent with cell wall compositional data (**Fig. 2**), S unit signals (**S**) were severely depleted in all *PMT*-deficient lines (**Fig. 3a,b**). Concurrently, diagnostic contours for naringenin (**N**) (Lam *et al*., 2017; Rencoret *et al*., 2022) and apigenin (**A**) (Lam *et al*., 2019; Rencoret *et al*., 2021) emerged in the *FNSII*- and *A3ʹH/C5ʹH*-deficient lignins, respectively (**Fig. 3a,c**). Although accumulating at substantially lower levels than native tricin levels in WT and *ospmt1/2* plants, the detection of these intermediates demonstrates that, in line with our earlier findings (Lam *et al*., 2017, 2019), upstream flavonoid precursors are partially integrated into the lignin polymer when downstream tricin biosynthesis is blocked.

**Fig. 3.**
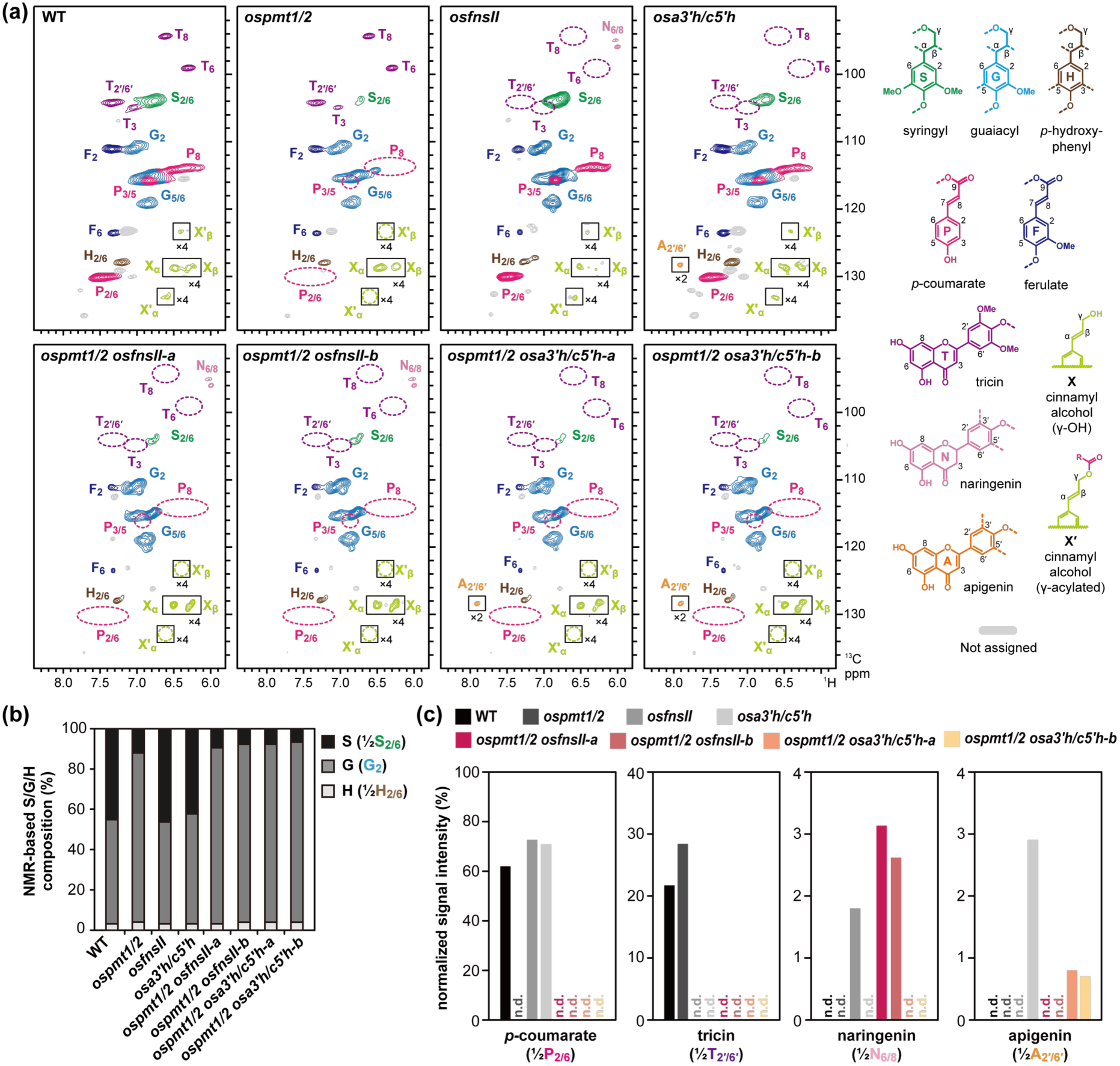
2D NMR analysis of the aromatic composition of dioxane/water-soluble lignin (DWL) samples extracted from *p*-coumarate- and tricin-deficient rice mutant cell walls. (a) Aromatic subregions of 2D HSQC NMR spectra of DWLs. DWLs were prepared from culm cell wall residue samples pooled from three biologically independent plants for each line. The contour coloration matches the lignin substructures shown. Boxes labeled ×2 and ×4 indicate regions with scale vertically enlarged for 2-fold and 4-fold, respectively. Peak assignments are listed in **Table S5**. (b and c) Volume integration analysis of major lignin aromatic units. NMR-based syringyl/guiacyl/*p*-hydroxyphenyl (S/G/H) unit composition (b) and signal intensities of *p*-coumarate (½**P_2/6_**), tricin (½**T_2ʹ/6ʹ_**) naringenin (½**N_6/8_**) and apigenin (½**A_2ʹ/6ʹ_**) units (c) are displayed. Volume integrals for the major lignin aromatic units are expressed as percentages relative to the total of S, G and H aromatic units (½**S_2/6_** + **G_2_** + ½**H_2/6_** = 100%). n.d., not detected; WT, wild type; *ospmt1/2*, *PMT*-deficient double-knockout line; *osfnsII*, *FNSII*-deficient single-knockout line; *osa3’h/c5’h*, *A3’H/C5’H*-deficient single-knockout line; *ospmt1/2 osfnsII-a/b*, *PMT*- and *FNSII*-deficient triple-knockout lines; *ospmt1/2 osa3’h/c5’h-a/b*, *PMT*- and *A3’H/C5’H*-deficient triple-knockout lines.

#### Lignin linkage patterns

The oxygenated aliphatic sub-regions (*δ*_C_/*δ*_H_, 90–52/6.0–2.5 ppm) of the 2D HSQC NMR spectra displayed signals attributed to the major lignin inter-unit linkages, such as β–O–4 (**I**), β–5 (**II**), β–β (**III** and **III’**), 5–5/β–O–4 (**IV**), and β–1 (**V**), as well as those attributed to the cinnamyl alcohol end-units (**X1** and **X1’**) (**Fig. 4a; Table S5**). The volume integration data of the major lignin inter-unit linkage and end-unit signals (percentages relative to **I_ɑ_** + **II_ɑ_** + ½**III_ɑ_** + ½**III′_β_** + **IV_ɑ_** + **V_ɑ_** = 100%) were used to estimate the differences in the lignin linkage patterns between the lignin samples (**Fig. 4b**). Due to the loss-of-function of the *PMT* genes, the signals from the tetrahydrofuran-type β–β (**IIIʹ**) units and γ-acylated cinnamyl alcohol end-units (**Xʹ**), both originating from the incorporation of monolignol *p*-coumarate conjugates (Lu & Ralph, 2002; Yamashita *et al*., 2020) were completely absent in the spectra of the DWL samples from all *PMT*-deficient lines (*ospmt1/2*, *ospmt1/2 osfnsII*, and *ospmt1/2 osa3ʹh/c5ʹh*); conversely, these diagnostic signals were readily detected in DWL samples from the WT, *osfnsII*, and *osa3ʹh/c5ʹh* lines (**Fig. 4a,b**). Furthermore, the flavanone C2–H2 correlations from naringenin (**N_2_**) (Lam *et al*., 2017; Rencoret *et al*., 2022) were clearly resolved in the spectra of all *FNSII*-deficient lines (*osfnsII* and *ospmt1/2 osfnsII*) (**Fig. 4a**), corroborating the participation of naringenin in lignification.

**Fig. 4.**
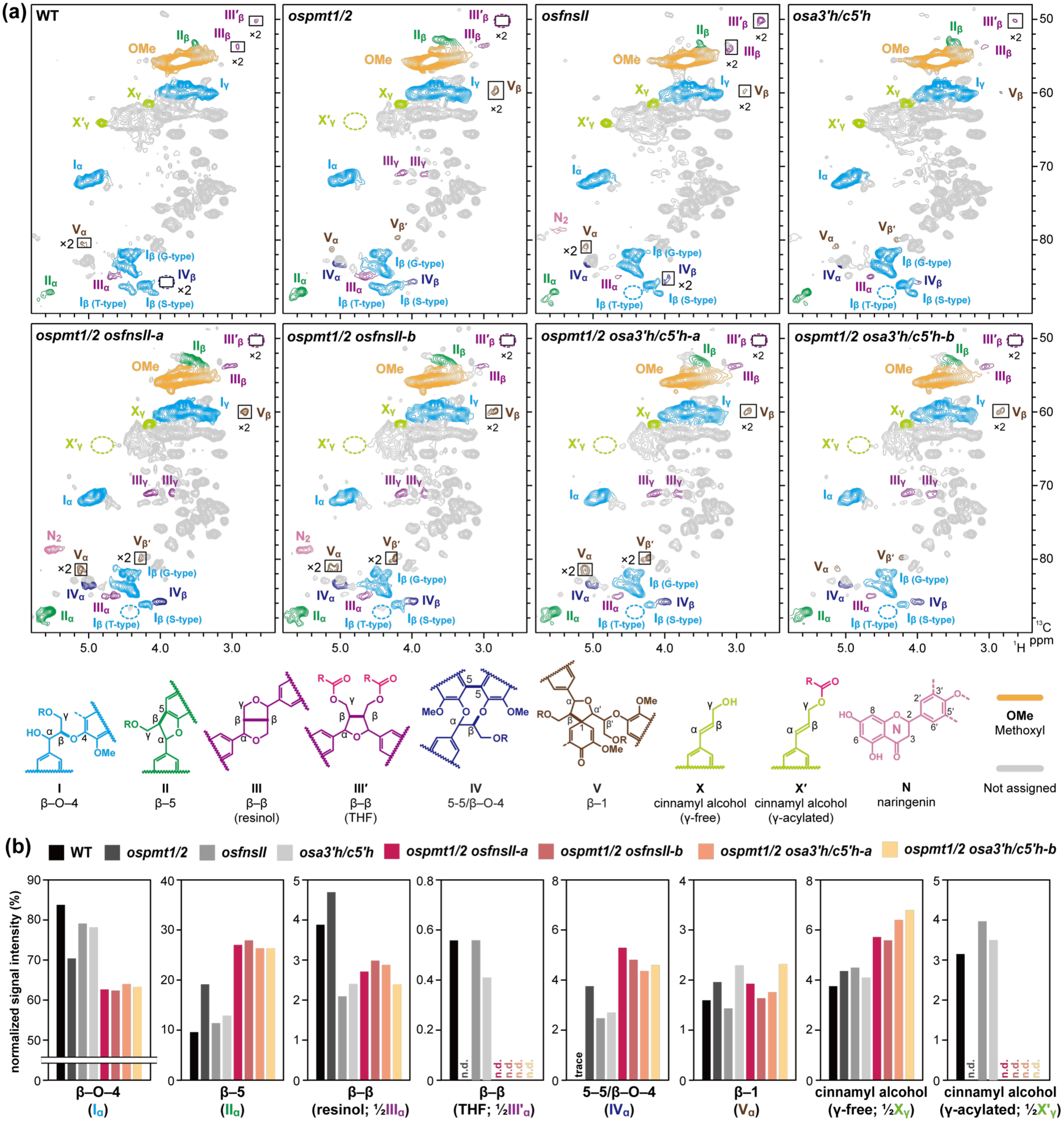
2D NMR analysis of the linkage distributions of dioxane/water-soluble lignin (DWL) samples extracted from *p*-coumarate- and tricin-deficient rice mutant cell walls. (a) Oxygenated aliphatic subregions of 2D HSQC NMR spectra of DWLs. DWLs were prepared from culm cell wall residue samples pooled from three biologically independent plants for each line. The contour coloration matches the lignin substructures shown. Boxes labeled ×2 indicate regions with scale vertically enlarged for 2-fold. Peak assignments are listed in **Table S5**. (b) Volume integration analysis of the major lignin intermonomeric linkage and end-unit types. Volume integrals are expressed as percentages relative to the total of the analyzed intermonomeric linkage types (**I_ɑ_** + **II_ɑ_** + ½**III_ɑ_** + ½**IIIʹ_β_** + **IV_β_** + **V_ɑ_** = 100%). n.d., not detected. WT, wild type; *ospmt1/2*, *PMT*-deficient double-knockout line; *osfnsII*, *FNSII*-deficient single-knockout line; *osa3’h/c5’h*, *A3’H/C5’H*-deficient single-knockout line; *ospmt1/2 osfnsII-a/b*, *PMT*- and *FNSII*-deficient triple-knockout lines; *ospmt1/2 osa3’h/c5’h-a/b*, *PMT*- and *A3’H/C5’H*-deficient triple-knockout lines.

Alongside the aforementioned diagnostic monomeric alterations, the *p*-coumarate- and tricin-deficient mutants exhibited substantial quantitative shifts in their overall lignin linkage distributions (**Fig. 4b**). Our analysis revealed that the independent disruptions of *p*-coumarate and tricin biosynthesis exert an additive reduction on β–O–4 linkage abundance. Relative to the WT control, normalized β– O–4 (**I**) signal intensities fell by ∼14% in *ospmt1/2* and 3–5% in *osfnsII* and *osa3ʹh/c5ʹh*, with a more severe reduction of ∼19–21% observed in the *ospmt1/2 osfnsII* and *ospmt1/2 osa3ʹh/c5ʹh* stacked mutants. This depletion in β–O–4 bonds was accompanied by a marked increase in the signals from β–5 (**II**) and 5–5/β–O–4 (**IV**) linkages, as well as those from cinnamyl alcohol end-units (**X1**). Meanwhile, β–β linkages (**III**) displayed divergent trajectories, rising in the *p*-coumarate-deficient *ospmt1/2* mutant but falling in the tricin-deficient *osfnsII* and *osa3ʹh/c5ʹh* mutants; consequently, these opposing effects largely offset each other in the *ospmt1/2 osfnsII* and *ospmt1/2 osa3ʹh/c5ʹh* stacked mutants, leaving overall β–β levels marginally affected. Meanwhile, β–1 (**V**) linkage frequency showed only minor variations across the *p*-coumarate- and tricin-deficient mutants.

#### Molecular weight distributions

To assess how the loss of *p*-coumarate and tricin units affects the lignin polymer size, we analyzed the DWL samples via GPC (**Fig. 5a**). The *p*-coumarate-deficient *ospmt1/2* mutant displayed number-average (*M*_n_) and weight-average (*M*_w_) molecular weights nearly identical to those of the WT control (*M*_n_, ∼2.8 vs. ∼2.7 kg mol^−1^; *M*_w_, ∼7.8 vs. ∼8.1 kg mol^−1^) (**Fig. 5b**). In contrast, every tricin-deficient line (*osfnsII*, *osa3ʹh/c5ʹh*, *ospmt1/2 osfnsII*, and *ospmt1/2 osa3ʹh/c5ʹh*) exhibited consistently elevated molecular weights (*M*_n_, ∼3.3–3.7 kg mol^−1^; *M*_w_, ∼9.1–10.0 kg mol^−1^) (**Fig. 5b**). These data indicate that the depletion of tricin rather than *p*-coumarate units contributes to an increase in overall molecular weight of isolated lignin.

**Fig. 5.**
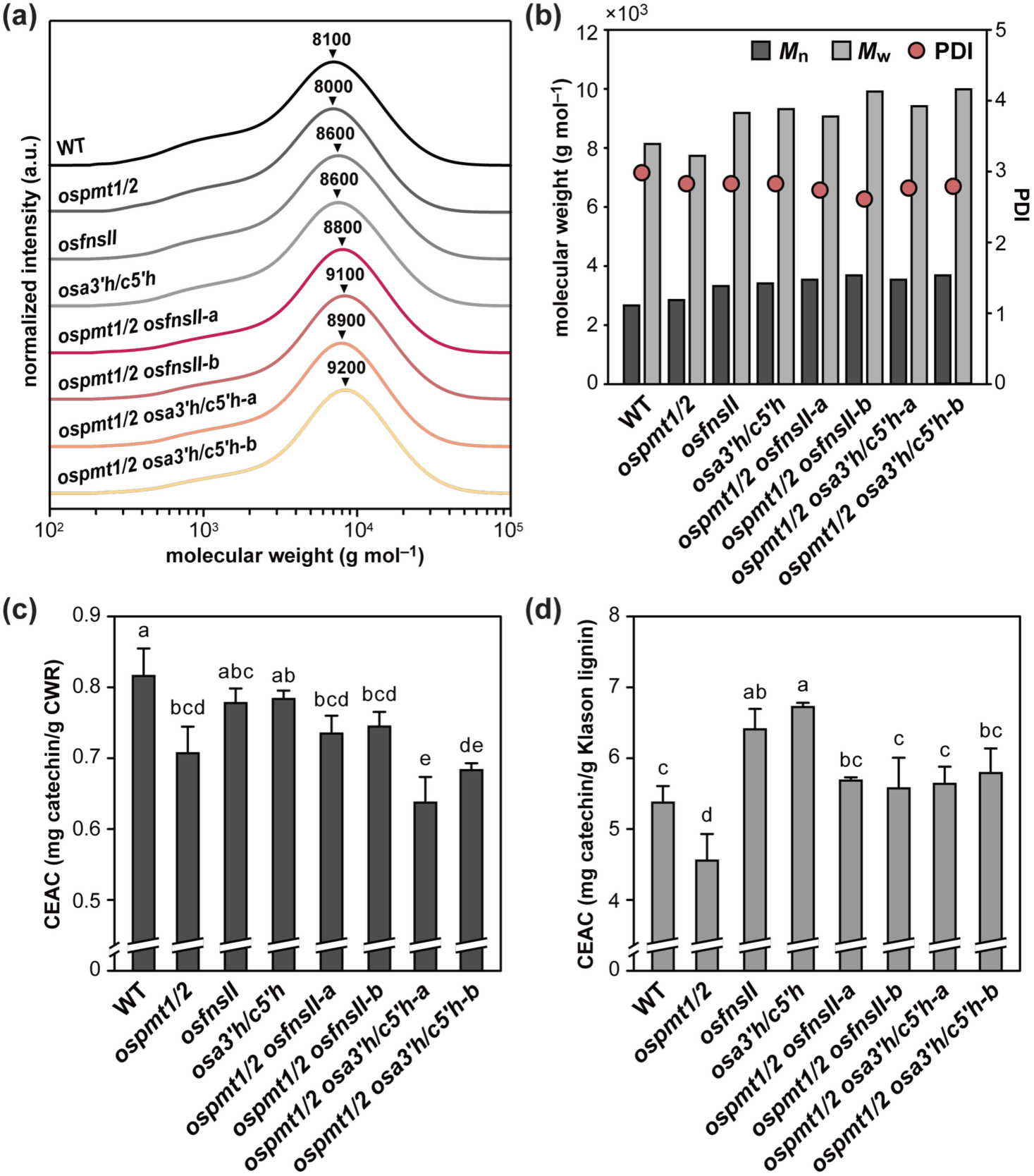
Analysis of lignin molecular weight and antioxidant activity in *p*-coumarate- and tricin-deficient rice mutant cell walls. (a and b) Gel permeation chromatography (GPC)-derived molecular weight distribution curves (a) and averaged molecular weight data (b) of dioxane/water-soluble lignin (DWL) samples extracted from rice culm cell wall residue (CWR) samples. DWLs were prepared from CWRs pooled from three biologically independent plants for each line. The molecular weights were calibrated using polystyrene standards. In (a), peak-top molecular weight (*M*_p_) is marked on each curve. *M*_n_, number-averaged molecular weight; *M*_w_, weight-averaged molecular weight; PDI, polydiversity index (*M*_w_/*M*_n_); a.u., arbitrary unit. (c and d) Antioxidant activity of culm CWR was evaluated using the 2,2-diphenyl-1-picrylhydrazyl (DPPH) assay and expressed as catechin equivalent antioxidant capacity (CEAC) per unit mass of CWR (c) or Klason lignin (d). The values are means ± standard deviation from individually analyzed plants (*n* = 3). Different letters indicate significant differences (one-way ANOVA with Tukey HSD test, *P* < 0.05). WT, wild type; *ospmt1/2*, *PMT*-deficient double-knockout line; *osfnsII*, *FNSII*-deficient single-knockout line; *osa3’h/c5’h*, *A3’H/C5’H*-deficient single-knockout line; *ospmt1/2 osfnsII-a/b*, *PMT*- and *FNSII*-deficient triple-knockout lines; *ospmt1/2 osa3’h/c5’h-a/b*, *PMT*- and *A3’H/C5’H*-deficient triple-knockout lines.

Overall, our NMR and GPC analyses of isolated lignins demonstrate that the elimination of *p*-coumarate and tricin units alters not only monomer composition, but also the inter-unit linkage distribution and macromolecular size of the lignin polymers. This reflects the distinct biochemical roles these grass-specific modification units play during lignin polymerization, as further discussed below.

### *p*-Coumarate rather than tricin drives the radical scavenging capacity of rice lignin

Because plant *p*-hydroxycinnamates and flavonoids, including free *p*-coumarate and tricin, are well-known antioxidants (Bors *et al*., 1997; Mohanlal *et al*., 2011; Boz, 2015), we investigated whether the *p*-coumarate and tricin modifications confer radical scavenging properties to rice lignin and cell walls. To test this, we evaluated the radical scavenging capacity of culm CWR samples from the *p*-coumarate- and tricin-deficient mutants using a modified DPPH assay optimized for insoluble polymeric substances (Serpen *et al*., 2007). Radical scavenging capacity, expressed as catechin equivalent antioxidant capacity (CEAC) per unit mass of CWR (Sawamura *et al*., 2017), was reduced across all mutants compared with the WT control, although this reduction lacked statistical significance in the *osfnsII* and *osa3ʹh/c5ʹh* lines (**Fig. 5c**). This overall decline could be attributed to decreases in *p*-coumarate and/or tricin units within lignin, and/or total lignin content in the mutant cell walls. When CEAC was normalized per unit mass of Klason lignin within each cell wall sample, the *p*-coumarate-deficient *ospmt1/2* mutant displayed reduced capacity, whereas the tricin-deficient *osfnsII* and *osa3ʹh/c5ʹh* mutants displayed increased capacity (**Fig. 5d**). Meanwhile, the *ospmt1/2 osfnsII* and *ospmt1/2 osa3ʹh/c5ʹh* stacked mutants exhibited intermediate capacities comparable to the WT control. Collectively, these data indicate that the incorporation of *p*-coumarate decoration units markedly enhances the radical scavenging properties of lignin, whereas tricin units conversely attenuate this capacity. The underlying molecular mechanisms governing these functional shifts are discussed below.

## Discussion

Grasses uniquely evolved *p*-coumarate and tricin units on their monolignol-derived lignin polymer backbones. Yet, the evolutionary, physiological, and biochemical significance of these grass-specific lignin modifications, as well as their impact on grass biomass valorization, remains poorly understood. To address this, we introduced stacked mutations to simultaneously block lignin *p*-coumaroylation and tricin biosynthesis in rice. Our chemical and NMR characterizations confirmed that the *ospmt1/2 osfnsII* and *ospmt1/2 osa3ʹh/c5ʹh* stacked mutants assemble lignin completely devoid of both modification units (**Fig. 2,3**). Collectively, these results demonstrate the successful engineering of a “eudicot-like” lignin polymer within a grass host, further corroborating the remarkable metabolic plasticity of cell wall lignification. While these *p*-coumarate- and tricin-deficient stacked mutants displayed minor growth reductions, their general viability confirms that *p*-coumarate and tricin units are largely dispensable for overall plant growth and development (**Table 1**). Whether these units confer selective functional advantages for cell wall properties and plant resilience under environmental stress remains an important question for future physiological studies. Here, we focused on elucidating their fundamental biochemical roles by synthesizing our in-depth lignin structural analysis with our cell wall antioxidant assays.

### Disrupting *p*-coumarate and tricin biosynthesis altered lignin monomer assembly

Beyond the complete elimination of *p*-coumarate and tricin units, the monolignol unit composition was substantially altered in the *ospmt1/2 osfnsII* and *ospmt1/2 osa3ʹh/c5ʹh* stacked mutants. Strikingly, the S/G unit ratio in these mutants dropped to ∼0.1 (based on NMR data of DWL samples), which is substantially lower than that of the WT control (∼0.5) (based on NMR data of DWL samples) (**Fig. 2,3**). Such a severe reduction in the S/G ratio is primarily attributable to the disruption of lignin *p*-coumaroylation, as an identically low S/G ratio was previously established in the *ospmt1/2* parent background (Lam *et al*., 2024b) (**Fig. 2,3**). Similar reductions in the S/G ratio upon disruptions of *PMT* genes have been also observed in maize (Marita *et al*., 2014; Oliveira *et al*., 2025) and *Brachypodium* (Petrick *et al*., 2014). Indeed, our recent work demonstrated that the PMT-initiated monolignol *p*-coumaroylation pathway is essential for S unit incorporation, functioning independently of the conventional S lignin biosynthetic pathway in rice and potentially across other grasses (Ji *et al*., 2025). Consequently, the lignin deposited in the *ospmt1/2 osfnsII* and *ospmt1/2 osa3ʹh/c5ʹh* mutants is predominantly composed of non-acylated G units, with only minimal remaining S and H units alongside the total absence of *p*-coumarate and tricin modification units. Such high monomeric homogeneity and marked G lignin enrichment provide an attractive feedstock for biorefinery valorization—including the catalytic production of high-value aromatic chemicals (Zakzeski *et al*., 2010; Ragauskas *et al*., 2014; Key & Bozell, 2016; Rinaldi *et al*., 2016; Oliveira *et al*., 2025) and advanced carbon materials (Li *et al*., 2018; Xu *et al*., 2024), alongside high-energy solid biofuel applications (Takeda *et al*., 2019b; Umezawa *et al*., 2018).

Our NMR analyses also corroborated the incorporation of naringenin and apigenin into lignin, in lieu of the canonical tricin monomer, upon the disruption of *FNSII* and *A3ʹH/C5ʹH*, respectively, albeit at substantially lower levels than native tricin in the WT controls (**Fig. 3**) (Lam *et al*., 2017, 2019a). Notably, naringenin incorporation increased in the *ospmt1/2 osfnsII* stacked mutants relative to the *osfnsII* single mutant, whereas conversely, apigenin incorporation decreased in the *ospmt1/2 osa3ʹh/c5ʹh* stacked mutants compared with its *osa3ʹh/c5ʹh* single-mutant counterpart (**Fig. 3**). These contrasting trends may imply that the radical polymerization preferences of naringenin and apigenin differ significantly depending on the presence of *p*-coumaroylated monolignols and/or the severe G-type monolignol enrichment characteristic of the *ospmt1/2* background. Validating these incorporation dynamics may provide mechanistic guidance for ongoing efforts to engineer non-canonical flavonoids and flavonolignins in bioenergy crops (Mahon *et al*., 2022).

### Elimination of *p*-coumarate and tricin decorating units alters lignin linkage distribution and molecular weight

Our in-depth structural analyses via 2D NMR and GPC revealed substantially altered lignin linkage distributions and molecular weights in the *p*-coumarate- and tricin-deficient mutants. This demonstrates that the elimination of these modification units impacts not only lignin monomer composition, but also the trajectory of their oxidative radical coupling during cell wall lignification. Crucially, knocking out lignin *p*-coumaroylation and tricin biosynthetic pathways results in an additive depletion of β–O–4 ether linkage abundance. This ether bond depletion was accompanied by a marked, compensatory increase in β–5 and 5–5/β–O–4 linkages alongside an elevated frequency of cinnamyl alcohol end-units (**Fig. 4**). The concomitant reduction in β–O–4 bonds and increase in β–5 and 5–5/β– O–4 linkages upon *PMT* disruption are likely correlated with the aforementioned reduction in the S/G ratio, as such shifts in linkage distribution are well-established consequences of G-unit enrichment over S-units, reflecting the distinct radical coupling preferences of G- and S-type monolignols (Marita *et al*., 1999; Anderson *et al*., 2015; Takeda *et al*., 2017, 2019a, 2019b; Ji *et al*., 2025). Conversely, the β–O–4 reduction observed in tricin-deficient lines can be attributed to the unique role of tricin as a starter unit or nucleation site during grass lignification (Lan *et al*., 2025). When initiating lignin polymerization, tricin cross-couples with monolignols or their *p*-coumarate conjugates exclusively via β–O–4-type (4ʹ–O–β) linkages (del Río *et al*., 2012; Lan *et al*., 2015, 2016; Elder *et al*., 2020; Berstis *et al*., 2021; Ralph *et al*., 2026). Thus, the genetic ablation of these tricin initiation sites inherently eliminates the associated β–*O*–4-type linkages, thereby reducing their overall frequency within the lignin polymer.

Notably, the elimination of *p*-coumarate versus tricin units exerts divergent effects on the molecular weight of isolated lignins (DWLs) (**Fig. 5**). Although we previously reported a notable molecular weight increase in *ospmt1/2* lignin (Lam *et al*., 2024b), the shift observed in the present study was less pronounced, exhibiting only a minor increase (by ∼100 g mol^−1^) in *M*_n_ compared with the WT control. Conversely, all tricin-deficient mutants examined in this study exhibited a significantly more pronounced increase in both *M*_n_ (by ∼600–1,000 g mol^−1^) and *M*_w_ (by ∼1,000–1,900 g mol^−1^). This suggests that the incorporation of tricin rather than *p*-coumarate substantially suppresses the overall molecular size of the lignin polymer (**Fig. 5**), although our molecular weight data from the GPC analysis of isolated, soluble lignins (DWLs) do not reflect the entire molecular weight of native cell wall lignin. Our observation supports the aforementioned role of tricin acting as an initiation site for lignin polymerization (Lan *et al*., 2025); in the absence of such initiation sites, dehydrogenative polymerization can yield longer macromolecular chains within a constant monomer pool size—a mechanism recently predicted by comparing lignins from diverse plants possessing or lacking tricin units (Ji *et al*., 2025).

### Divergent roles of *p*-coumarate and tricin in dictating lignin antioxidant capacity

The antioxidant activity of lignin has been extensively investigated, driven both by its physiological relevance in protecting plant cell walls against oxidative stress and by its potential for high-value material applications (Dizhbite *et al*., 2004; Lu *et al*., 2022; Sadeghifar & Ragauskas, 2025; Yang *et al*., 2025; Yu *et al*., 2025). Because both *p*-coumarate and tricin are well known to exhibit strong antioxidant activity in their free forms (Bors *et al*., 1997; Korkina, 2007; Strandås *et al*., 2008; Mohanlal *et al*., 2011; Ajitha *et al*., 2012; Boz, 2015), we investigated whether their integration alters the antioxidant properties of grass lignin, and consequently, whether cell walls assembled by our *p*-coumarate- and tricin-deficient mutants display altered antioxidant capacities. Our comparative analysis of DPPH radical scavenging assays across the mutant lines demonstrated that the incorporation of *p*-coumarate decorating units markedly enhances the radical scavenging properties of rice lignin. In contrast, and somewhat counterintuitively, tricin units attenuate this capacity (**Fig. 5**).

Free phenolic hydroxyl (OH) groups primarily drive the radical scavenging activity of phenolic antioxidants, including lignin (Sawamura *et al*., 2017; Li *et al*., 2024; Sadeghifar & Ragauskas, 2025) and *p*-coumarates (Cai *et al*., 2006; Strandås *et al*., 2008). Because ester-linked *p*-coumarate pendant units retain these free phenolic OH groups, they directly impart radical scavenging capacity to the assembled lignin polymer. Conversely, in some flavonoid types including tricin, radical scavenging capacity is primarily governed by the B-ring phenolic OH groups, whereas the *meta*-positioned A-ring OH groups themselves exhibit negligible radical scavenging capacity (Rice-Evans *et al*., 1996; Arora *et al*., 1998; Cai *et al*., 2006; Thavasi *et al*., 2009). Indeed, tricin’s DPPH scavenging activity drops dramatically when its B-ring OH group is blocked (Ajitha *et al*., 2012). Because tricin integrates into the lignin backbone exclusively via 4ʹ–O–β ether linkages that mask this essential B-ring OH group (del Río *et al*., 2012; Lan *et al*., 2015, 2016; Elder *et al*., 2020; Berstis *et al*., 2021; Ralph *et al*., 2026), it cannot confer radical scavenging capacity to the polymer matrix. Furthermore, because eliminating tricin units increases the average lignin polymer chain length (**Fig. 5**), it proportionally reduces the frequency of phenolic OH end-groups across lignin macromolecule, further attenuating its intrinsic radical scavenging capacity.

Because a plant’s antioxidant capacity correlates with its ability to scavenge reactive oxygen species (ROS) generated during biotic and abiotic stress (Hasanuzzaman *et al*., 2012; Nakabayashi *et al*., 2014; Das & Roychoudhury, 2014; Kerchev & Van Breusegem, 2022), *p*-coumarate decorations in grass lignin may enhance environmental stress tolerance by contributing to ROS detoxification. Furthermore, because recent biorefinery studies indicate that lignin-bound tricin substantially improves the UV-screening capacity of lignin-derived materials (Zhang *et al*., 2025), it is intriguing to hypothesize that tricin also confer UV resilience to cell wall lignin *in planta*. The physiological significance of grass lignin modifications can be systematically elucidated by assessing stress performance in the *p*-coumarate- and tricin-deficient mutants developed in this study, alongside complementary overexpression lines overproducing these lignin modification units. These insights will guide targeted breeding and metabolic engineering strategies in grasses to co-optimize plant growth, stress resilience, and food and biomass production.

## Acknowledgements

We thank Prof. Keishi Osakabe (Tokushima University) and Prof. Yuriko Osakabe (Institute of Science Tokyo) for providing the pMgPoef4_129-2A-GFP vector and for their helpful suggestions regarding the genome editing experiments, as well as Prof. Hironori Kaji and Ms. Ayaka Maeno (Kyoto University) for their assistance with the NMR analysis. This work was supported in part by grants from the Japan Society for the Promotion of Science (JSPS KAKENHI Grant Nos. JP20H03044 and JP24K01827). This study was partially conducted using the greenhouse and GC-MS facilities of the Development and Assessment of Sustainable Humanosphere/Forest Biomass Analytical System (DASH/FBAS, Research Institute for Sustainable Humanosphere, Kyoto University) and the NMR spectrometer at the International Joint Usage/Research Center (iJURC, Institute for Chemical Research, Kyoto University).

## Author contributions

LPYL, TU, and YT conceptualized and designed the study. LPYL, SY, and YT conducted the experiments and analyzed the data. LPYL and YT wrote the original draft of the manuscript, which was reviewed and edited by SY and TU.

## Data availability

The authors confirm that the data supporting the findings of this study are available within the article and its supplementary materials. The accession numbers for *PMT*, *FNSII*, and *A3ʹH/C5ʹH* genes studied in this study are listed in the Materials and Methods section.

## Supporting Information

The following Supporting Information is available for this article:

**Fig. S1** Mutation patterns of *OsPMT1* and *OsPMT2* in *PMT*-deficient rice mutants.

**Fig. S2** Predicted effects of mutations on *OsFNSII* in *FNSII*-deficient rice mutants.

**Fig. S3** Predicted effects of mutations on *OsA3ʹH/C5ʹH* in *A3ʹH/C5ʹH*-deficient mutants.

**Fig. S4** 2D HSQC NMR spectra of rice cell walls.

**Fig. S5** Volume integration analysis of 2D HSQC NMR spectra of rice cell walls.

**Table S1** Primers and oligonucleotides used in this study.

**Table S2** Off-target analysis of genome-edited rice mutants.

**Table S3** Neutral sugar analysis of rice cell walls.

**Table S4** Peak assignments in 2D HSQC NMR spectra of rice cell walls.

**Table S5** Peak assignments in 2D HSQC NMR spectra of dioxane/water-soluble lignins.

**Supplementary References**

