## Supporting Information for "Engineering eudicot-like lignin in rice via targeted disruption of grass-specific lignin modification pathways"

The following Supporting Information is available for this article:

**Fig. S1.** Mutation patterns of *OsPMT1* and *OsPMT2* in *PMT*-deficient rice mutants.

**Fig. S2.** Predicted effects of mutations on *OsFNSII* in *FNSII*-deficient rice mutants.

**Fig. S3.** Predicted effects of mutations on *OsA3'H/C5'H* in *A3'H/C5'H*-deficient mutants.

**Fig. S4.** 2D HSQC NMR spectra of rice cell walls.

**Fig. S5.** Volume integration analysis of 2D HSQC NMR spectra of rice cell walls.

**Table S1.** Primers and oligonucleotides used in this study.

**Table S2.** Off-target analysis of genome-edited rice mutants.

**Table S3.** Neutral sugar analysis of rice cell walls.

**Table S4**. Peak assignments in 2D HSQC NMR spectra of rice cell walls.

**Table S5.** Peak assignments in 2D HSQC NMR spectra of dioxane/water-soluble lignins.

**Supplementary References**

**Fig. S1** Mutation patterns of *OsPMT1* and *OsPMT2* in *PMT*-deficient rice mutants. The *ospmt1/2* double-knockout line was originally developed by Lam et al. (2024). WT, wild type; *ospmt1/2*, *PMT*-deficient double-knockout line; *ospmt1/2 osfnsII-a/b*, *PMT*- and *FNSII*-deficient triple-knockout lines; *ospmt1/2 osa3'h/c5'h-a/b*, *PMT*- and *A3'H/C5'H*-deficient triple-knockout lines.

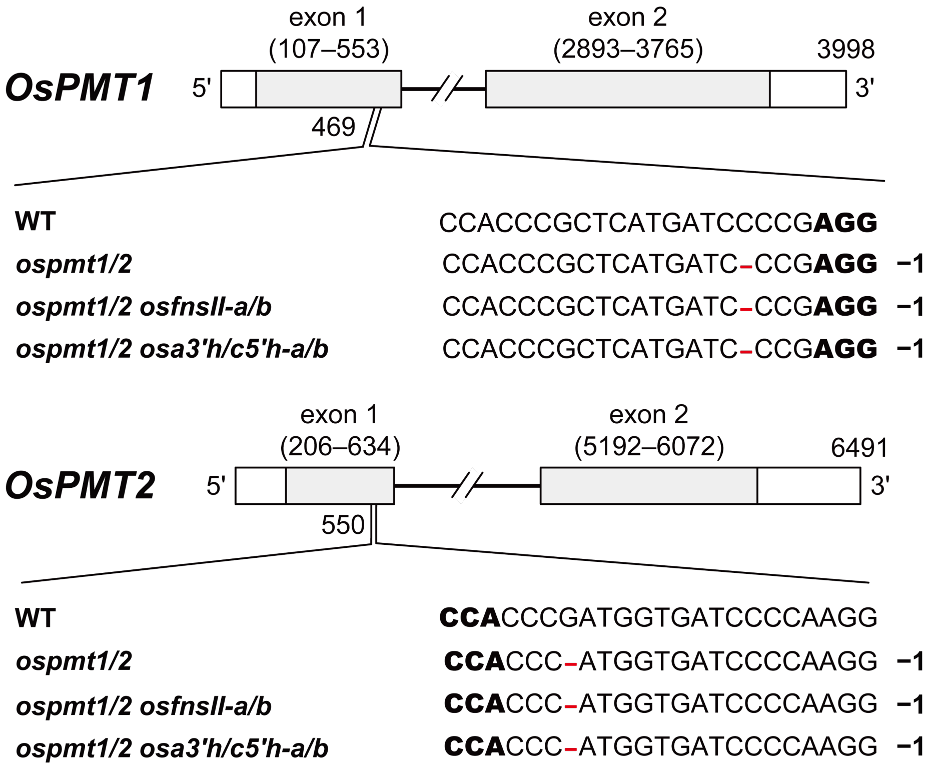

**Fig. S2** Predicted effects of mutations on *OsFNSII* in *FNSII*-deficient rice mutants. Protein sequence alignment of the wild-type and mutated OsFNSII proteins in the *osfnsII*, *ospmt1/2 osfnsII-a*, and *ospmt1/2 osfnsII-b* mutants. Asterisks indicate stop codons. The first premature stop codon is shown in red. The blue text indicates amino acid sequence differences from the wild-type OsFNSII protein, assuming translation starts from the native start codon. WT, wild type; *osfnsII*, *FNSII*-deficient single-knockout line; *ospmt1/2 osfnsII-a/b*, *PMT*- and *FNSII*-deficient triple-knockout lines.

```
>FNSII
                                                              hinge region
WT                   MASLMEVQVPLLGMGTTMGALALALVVVVVVHVAVNAFGRRRLPPSPASLPVIGHLHLLR   60
osfnsII              MASLMEVQVPLLGMGTTMGALALALVVVVVVHVAVNAFGRRRLPPSPASLPVIGHLHLLR   60
ospmt1/2 osfnsII-a   MASLMEVQVPLLGMGTTMGALALALVVVVVVHVAVNAFGRRRLPPSPASLPVIGHLHLLR   60
ospmt1/2 osfnsII-b   MASLMEVQVPLLGMGTTMGALALALVVVVVVHVAVNAFGRRRLPPSPASLPVIGHLHLLR   60

WT                   PPVHRTFHELAARLGPLMHVRLGSTHCVVASSAEVAAELIRSHEAKISERPLTAVARQFA  120
osfnsII              PPVHRTFHELAARLGPLMHVRLGSTHCVVASSAEVAAELIRSHEAKISERPLTAVARQFA  120
ospmt1/2 osfnsII-a   PPVHRTFHELAARLGPLMHVRLGSTHCVVASSAEVAAELIRSHEAKISERPLTAVARQFA  120
ospmt1/2 osfnsII-b   PPVHRTFHELAARLGPLMHVRLGSTHCVVASSAEVAAELIRSHEAKISERPLTAVARQFA  120

WT                   YESAGFAFAPYSPHWRFMKRLCMSELLGPRTVEQLRPVRRAGLVSLLRHVLSQPEAEAVD  180
osfnsII              YESAGFAFAPYSPHWRFMKRLCMSELLGPRTVEQLRPVRRAGLVSLLRHVLSQPEAEAVD  180
ospmt1/2 osfnsII-a   YESAGFAFAPYSPHWRFMKRLCMSELLGPRTVEQLRPVRRAGLVSLLRHVLSQPEAEAVD  180
ospmt1/2 osfnsII-b   YESAGFAFAPYSPHWRFMKRLCMSELLGPRTVEQLRPVRRAGLVSLLRHVLSQPEAEAVD  180

WT                   LTRELIRMSNTSIIRMAASTVPSSVTEEAQELVKVVAELVGAFNADDYIALCRGWDLQGL  240
osfnsII              LTRELIRMSNTSIIRMAASTVPSSVTEEAQELVKVVAELVGAFNADDYIALCRGWDLQGL  240
ospmt1/2 osfnsII-a   LTRELIRMSNTSIIRMAASTVPSSVTEEAQELVKVVAELVGAFNADDYIALCRGWDLQGL  240
ospmt1/2 osfnsII-b   LTRELIRMSNTSIIRMAASTVPSSVTEEAQELVKVVAELVGAFNADDYIALCRGWDLQGL  240

WT                   GRRAADVHKRFDALLEEMIRHKEEARMRKKTDTDVGSKDLLDILLDKAEDGAAEVKLTRD  300
osfnsII              GRRAADVHKRFDALLEEMIRHKEEARMRKKTDTDVGSKDLLDILLDKAEDGAAEVKLTRD  300
ospmt1/2 osfnsII-a   GRRAADVHKRFDALLEEMIRHKEEARMRKKTDTDVGSKDLLDILLDKAEDGAAEVKLTRD  300
ospmt1/2 osfnsII-b   GRRAADVHKRFDALLEEMIRHKEEARMRKKTDTDVGSKDLLDILLDKAEDGAAEVKLTRD  300
                           oxygen binding pocket
WT                   NIKAFIIDVVTAGSDTSAAM--VEWMVAELMNHPEALRKVREEIEA---------VVGRD  349
osfnsII              NIKAFIIDVVTAGSDNFGRHGGV---DGGGADEP---------------PGGPAQGAGGD  342
ospmt1/2 osfnsII-a   NIKAFIIDVVTAG--SFGRHGGV---DGGGADEP---------------PGGPAQGAGGD  340
ospmt1/2 osfnsII-b   NIKAFIIDVVTAGSDTSRPPWWSGWWRS**TTRRPCARCGRRSRRWWGGTGSPARGTCRD  358
                                     ExxR motif
WT                   -RIAGEGDLPRLPYLQAAYKETLRLRPAAPIAHRQSTEEIQIRGFRVPAQTAVFINVWAI  408
osfnsII              RGGGGAGQDR------------RR---GGPAETAVSAGGV-QGDAAVEAGGADR-AQAVD  385
ospmt1/2 osfnsII-a   RGGGGAGQDR------------RR---GGPAETAVSAGGV-QGDAAVEAGGADR-AQAVD  383
ospmt1/2 osfnsII-b   CRICR-RRTR------------RR---CG*GRRRRSRTGSRRRRSRSEGSGCRR-RRRCS  400

WT                   GRDPAY-----------WEEPLEFRPERFLAGGGGE--GVE----------------PRG  439
osfnsII              GGDPDPRVQGAGADGGVHQRVGHRARPGVLGGAAGVQAGAVPRRRRRRGRGAARAALPVH  445
ospmt1/2 osfnsII-a   GGDPDPRVQGAGADGGVHQRVGHRARPGVLGGAAGVQAGAVPRRRRRRGRGAARAALPVH  443
ospmt1/2 osfnsII-b   --------STCGPS-GETRRTGRS---RWSSGRSGS--------------------SPAA  428
                        heme binding domain
WT                   QHFQFMPFGSGRRGCPGMGLALQSVPAVVAALLQCFDWQCMD--NKLIDMEEADGLVCAR  497
osfnsII              AVRERPARVPRDGACAAVGAGGGGGAAAVLRLAVHGQ*VDRH-----GG---------GR  490
ospmt1/2 osfnsII-a   AVRERPARVPRDGACAAVGAGGGGGAAAVLRLAVHGQ*VDRH-----GG---------GR  488
ospmt1/2 osfnsII-b   AARAWSRAGSTSSSCRSG-AAGAGAPGWGLRCSRCRRWWRRCCSASIGSAWTIS**TWRR  485

WT                   KHR-----LLLHAHPRLHPFPPLL*---------   516
osfnsII              R---PGLRSEASPPPPRPPAPPPFPAAPL-----   516
ospmt1/2 osfnsII-a   R---PGLRSEASPPPPRPPAPPPFPAAPL-----   514
ospmt1/2 osfnsII-b   QTAWFALGSIASSST-------PTRASTLSRRSS   512
```

**Fig. S3** Predicted effects of mutations on *OsA3'H/C5'H* in *A3'H/C5'H*-deficient mutants. Protein sequence alignment of the wild-type and mutated OsA3'H/C5'H proteins in the *osa3'h/c5'h*, *ospmt1/2 osa3'h/c5'h-a*, and *ospmt1/2 osa3'h/c5'h-b* mutants. Asterisks indicate stop codons. The first premature stop codon is shown in red. The blue text indicates amino acid sequence differences from the wild-type OsA3'H/C5'H protein, assuming translation starts from the native start codon. WT, wild type; *osa3'h/c5'h*, *A3'H/C5'H*-deficient single-knockout line; *ospmt1/2 osa3'h/c5'h-a/b*, *PMT*- and *A3'H/C5'H*-deficient triple-knockout lines.

```
>A3'H/C5'H
                                                                   hinge region
WT                        MEVAAMEISTSLLLTTVALSVIVCYALVFSRAGKARAPLPLPPGPRGWPVLGNLPQLGGK   60
oa3'hc5'h                 MEVAAMEISTSLLLTTVALSVIVCYALVFSRAGKARAPLPLPPGPRGWPVLGNLPQLGGK   60
ospmt1/2 oa3'hc5'h-a      MEVAAMEISTSLLLTTVALSVIVCYALVFSRAGKARAPLPLPPGPRGWPVLGNLPQLGGK   60
ospmt1/2 oa3'hc5'h-b      MEVAAMEISTSLLLTTVALSVIVCYALVFSRAGKARAPLPLPPGPRGWPVLGNLPQLGGK   60

WT                        THQTLHEMTKVYGPLIRLRFGSSDVVVAGSAPVAAQFLRTHDANFSSRPRNSGGEHMAYN   120
oa3'hc5'h                 THQTLHEMTKVYGPLIRLRFGSSDVVVAGSAPVAAQFLRTHDANFSSRPRNSGGEHMAYN   120
ospmt1/2 oa3'hc5'h-a      THQTLHEMTKVYGPLIRLRFGSSDVVVAGSAPVAAQFLRTHDANFSSRPRNSGGEHMAYN   120
ospmt1/2 oa3'hc5'h-b      THQTLHEMTKVYGPLIRLRFGSSDVVVAGSAPVAAQFLRTHDANFSSRPRNSGGEHMAYN   120

WT                        GRDVVFGPYGPRWRAMRKICAVNLFSARALDDLRAFREREAVLMVRSLAEASAAPGSSSP   180
oa3'hc5'h                 GRDVVFGPYGPRWRAMRKICAVNLFSARALDDLRAFREREAVLMVRSLAEASAAPGSSSP   180
ospmt1/2 oa3'hc5'h-a      GRDVVFGPYGPRWRAMRKICAVNLFSARALDDLRAFREREAVLMVRSLAEASAAPGSSSP   180
ospmt1/2 oa3'hc5'h-b      GRDVVFGPYGPRWRAMRKICAVNLFSARALDDLRAFREREAVLMVRSLAEASAAPGSSSP   180

WT                        AAVVLGKEVNVCTTNALSRAAVGRRVFAAGAGEGAREFKEIVLEVMEVGGVLNVGDFVPA   240
oa3'hc5'h                 AAVVLGKEVNVCTTNALSRAAVGRRVFAAGAGEGAREFKEIVLEVMEVGGVLNVGDFVPA   240
ospmt1/2 oa3'hc5'h-a      AAVVLGKEVNVCTTNALSRAAVGRRVFAAGAGEGAREFKEIVLEVMEVGGVLNVGDFVPA   240
ospmt1/2 oa3'hc5'h-b      AAVVLGKEVNVCTTNALSRAAVGRRVFAAGAGEGAREFKEIVLEVMEVGGVLNVGDFVPA   240

WT                        LRWLDPQGVVARMKKLHRRFDDMMNAIIAERRAGSLLKPTDSREEGKDLLGLLLAMVQEQ   300
oa3'hc5'h                 LRWLDPQGVVARMKKLHRRFDDMMNAIIAERRAGSLLKPTDSREEGKDLLGLLLAMVQEQ   300
ospmt1/2 oa3'hc5'h-a      LRWLDPQGVVARMKKLHRRFDDMMNAIIAERRAGSLLKPTDSREEGKDLLGLLLAMVQEQ   300
ospmt1/2 oa3'hc5'h-b      LRWLDPQGVVARMKKLHRRFDDMMNAIIAERRAGSLLKPTDSREEGKDLLGLLLAMVQEQ   300
                                                  oxygen binding pocket
WT                        EWLAAGEDDRITDTEIKALILNLFVAGTDTTSTIVEWTMAELIRHPDILKHAQEELDVVV   360
oa3'hc5'h                 EWLAAGEDDRITDTEIKALILNLFVAGTDTTSTIVEWTMAELIRHPDILKHKRS*MLLWV   359
ospmt1/2 oa3'hc5'h-a      EWLAAGEDDRITDTEIKALILNLFVAGTDTTSTIVEWTMAELIRHPDILKHKRS*MLLWV   359
ospmt1/2 oa3'hc5'h-b      EWLAAGEDDRITDTEIKALILNLFVAGTDTTSTIVEWTMAELIRHPDILKQRGARCCCGS   360
                                                                               ExxR motif
WT                        --GRD--RLLSESDLSHL-TFFHAI-I-------------------------KETFRLHP   389
oa3'hc5'h                 VIGSSQSRIYHISPSSMLSSRRHSVYIHQHRSRCHAWHLRSVRSQATVSPRVQSCWSMCG   419
ospmt1/2 oa3'hc5'h-a      VIGSSQSRIYHISPSSMLSSRRHSVYIHQHRSRCHAWHLRSVRSQATVSPRVQSCWSMCG   419
ospmt1/2 oa3'hc5'h-b      --**A--PLRVGSITSHLLPCYHQGDIPST----------------SINTALAATHGI*G   397

WT                        STPLSLPRMASEECEIAGYRIPKGAE----------------LLVNVWGIA---------   424
oa3'hc5'h                 GSPVTQPYGLTH*-----STSPLGSSPVGRTLMWMSREMISDLYHSVQGEGYAPASVGAC   473
ospmt1/2 oa3'hc5'h-a      GSPVTQPYGLTH*-----STSPLGSSPVGRTLMWMSREMISDLYHSVQGEGYAPASVGAC   473
ospmt1/2 oa3'hc5'h-b      V*DRRLP--YPQGCRVAGQCVGDRP*PSHMA-----------*PTRVQ-AL---------   431

WT                        ----RDPAIWPDPL------------------EYKPSRFLPGGTHTDVDVKGNDFGLIPF   462
oa3'hc5'h                 GWSP*QRPRWCMHSTGSYQRTRRQ---------------------------TSS---IWM   502
ospmt1/2 oa3'hc5'h-a      GWSP*QRPRWCMHSTGSYQRTRRQ---------------------------TSS---IWM   502
ospmt1/2 oa3'hc5'h-b      ----SVPPRWDAH*CGCQGK*FRTYTIRCRAKDMRRPQLGPADGHHDSGHAGACIRL---   482
                      heme binding domain
WT                        GAGRRICAGLSWGLRMVTMTAATLVHAFDWQLPADQTPDKLNMDEAFTLLLQRAEPLVVH   522
oa3'hc5'h                 RRLPSCCKGQS-----------------HWW---------------FTRYQGFSHPLTIL   530
ospmt1/2 oa3'hc5'h-a      RRLPSCCKGQS-----------------HWW---------------FTRYQGFSHPLTIL   530
ospmt1/2 oa3'hc5'h-b      --------------------------------AATSGPDARQAQYG*GVYPPAAKGRAIG   509

WT                        PVPR--L--LPSAYNIA*    535
oa3'hc5'h                 H-----------------    531
ospmt1/2 oa3'hc5'h-a      H-----------------    531
ospmt1/2 oa3'hc5'h-b      GSPGTKASPIRLQYCI--    525
```

**Fig. S4** 2D HSQC NMR spectra of whole cell wall samples from *p*-coumarate- and tricin-deficient rice mutants. Ball-milled rice culm cell walls (pooled from three biologically independent plants for each line) were subjected to the gel-state cell wall NMR measurements (Kim and Ralph 2009; Mansfield et al., 2012). Boxes labeled ×2 and ×4 indicate regions with scale vertically enlarged for 2- and 4-fold, respectively. Peak assignments are listed in **Table S4**. WT, wild type; *ospmt1/2*, *PMT*-deficient double-knockout line; *osfnsII*, *FNSII*-deficient single-knockout line; *osa3'h/c5'h*, *A3'H/C5'H*-deficient single-knockout line; *ospmt1/2 osfnsII-a/b*, *PMT*- and *FNSII*-deficient triple-knockout lines; *ospmt1/2 osa3'h/c5'h-a/b*, *PMT*- and *A3'H/C5'H*-deficient triple-knockout lines.

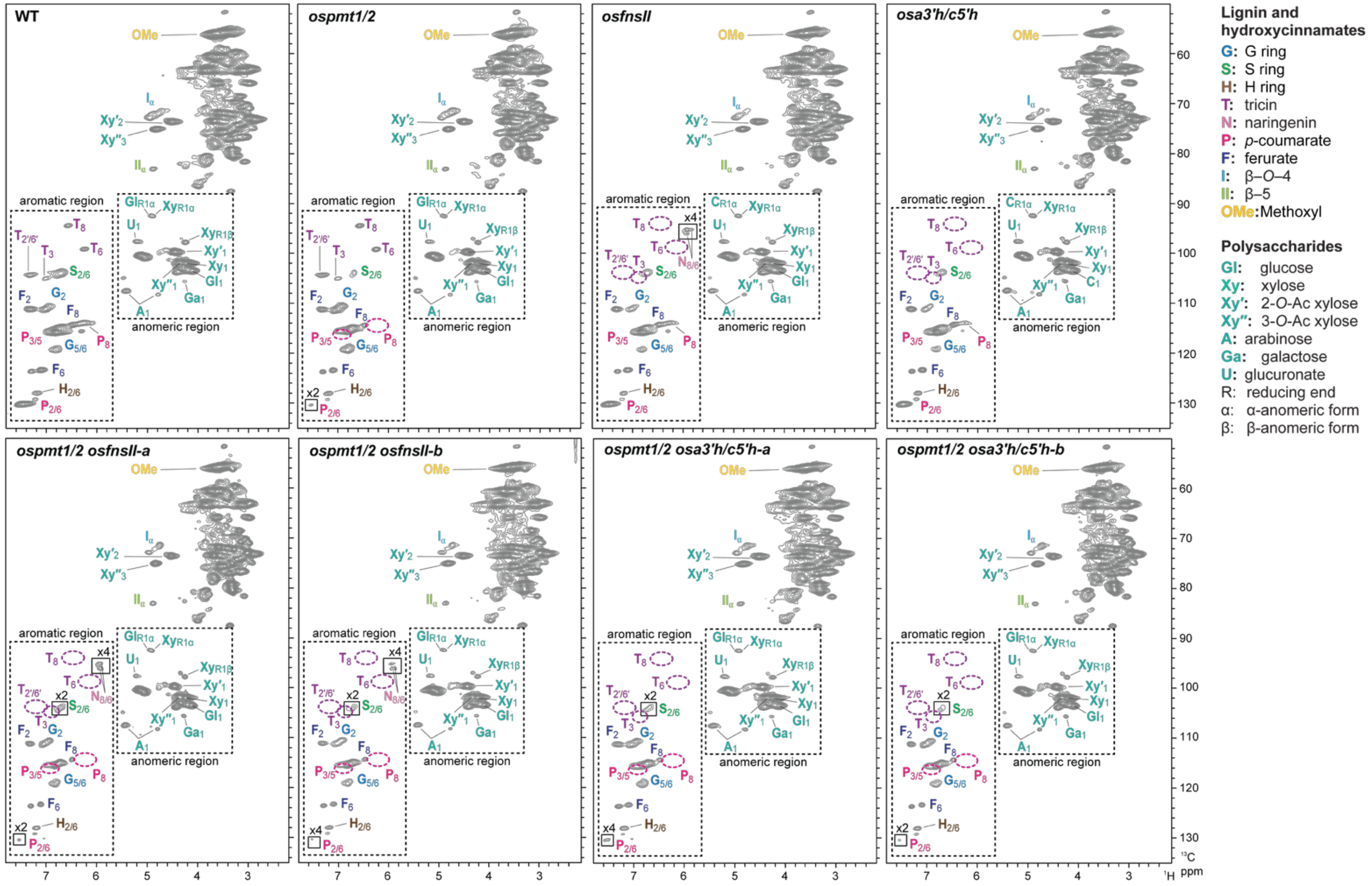

**Fig. S5** Volume integration analysis of 2D HSQC NMR spectra of whole cell walls from *p*-coumarate- and tricin-deficient rice mutants. Normalized signal intensities of major lignin, hydroxycinnamate, and polysaccharide units expressed as percentages of the sum of the listed signals ($\frac{1}{2}\mathbf{S_{2/6}} + \mathbf{G2} + \frac{1}{2}\mathbf{H_{2/6}} + \frac{1}{2}\mathbf{P_{2/6}} + \mathbf{F_2} + \frac{1}{2}\mathbf{T_{2'/6'}} + \mathbf{Gl_1} + \mathbf{X_1} + \mathbf{X'_1} + \mathbf{X''_1} + \mathbf{A_1} + \mathbf{GA_1} + \mathbf{U_1} = 100\%$) (Yamamoto et al., 2024; Ji et al., 2025). NMR analysis was conducted for culm CWR samples pooled from three biologically independent plants for each line. Peak assignments are listed in **Table S4**. n.d., not detected; WT, wild type; *ospmt1/2*, *PMT*-deficient double-knockout line; *osfnsII*, *FNSII*-deficient single-knockout line; *osa3'h/c5'h*, *A3'H/C5'H*-deficient single-knockout line; *ospmt1/2 osfnsII-a/b*, *PMT*- and *FNSII*-deficient triple-knockout lines; *ospmt1/2 osa3'h/c5'h-a/b*, *PMT*- and *A3'H/C5'H*-deficient triple-knockout lines.

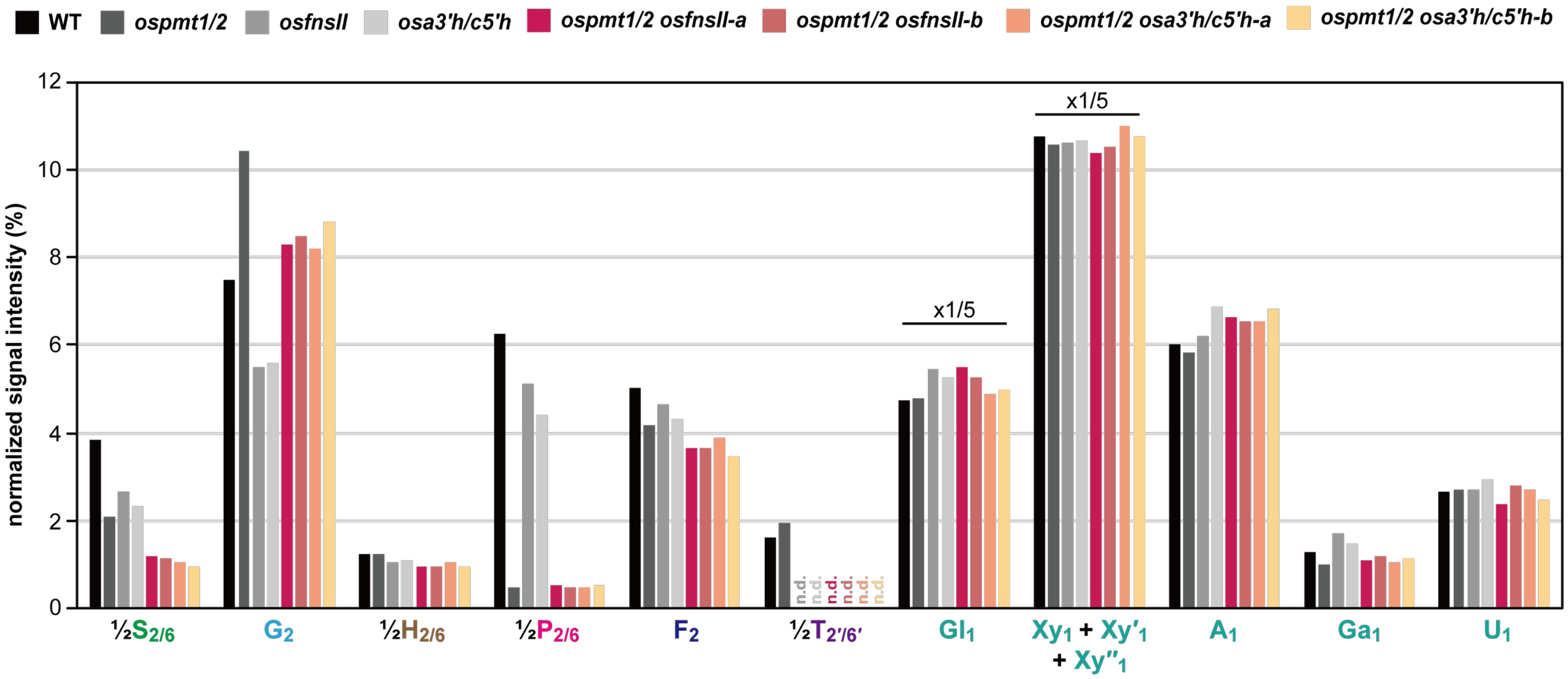

**Table S1** Primers and oligonucleotides used in this study.

| Primer name | Sequence (from 5´ to 3´) | Target gene/site | Purposes |
|---|---|---|---|
| LY92 | TGCATACTGCCGGGTCCGACACTT | *OsFNSII* | Generation of the *osfnsII* and *ospmt1/2 osfnsII* mutants |
| LY93 | AAACAAGTGTCGGACCCGGCAGTA | | |
| LY94 | TGCATATCCTCAAGCACGCCCAAG | *OsA3'H/C5'H* | Generation of the *osa3'h/c5'h* and *ospmt1/2 osa3'h/c5'h* mutants |
| LY95 | AAACCTTGGGCGTGCTTGAGGATA | | |
| LY100 | TGATCAGGCACAAGGAGGAG | *OsFNSII* | Genotyping of the *osfnsII* and *ospmt1/2 osfnsII* mutants |
| LY101 | ACCCTGAACCCTCGGATCT | | |
| LY102 | TTGAAAACATTGCCACATGAATG | *OsA3'H/C5'H* | Genotyping of the *osa3'h/c5'h* and *ospmt1/2 osa3'h/c5'h* mutants |
| LY103 | TTCCCTTGACATCCACATCA | | |
| LY105 | CACCCTTGATGTAGTCGGAGT | sgRNA-b-1[a] | Off-target analysis of the *osa3'h/c5'h* and *ospmt1/2 osa3'h/c5'h* mutants |
| LY106 | TCTCAAAGCCCTAGCTCCAA | | |
| LY107 | TGCCATGTTGGGGTCTATCT | sgRNA-b-2[a] | Off-target analysis of the *osa3'h/c5'h* and *ospmt1/2 osa3'h/c5'h* mutants |
| LY108 | CATTCGCAAAACGAGAACAA | | |
| LY109 | ACACGAAGAAGGCAATGGAG | sgRNA-b-3[a] | Off-target analysis of the *osa3'h/c5'h* and *ospmt1/2 osa3'h/c5'h* mutants |
| LY110 | GTTACAGCGTGGAACGACCT | | |

[a]Information of the potential off-target sites can be found in **Table S2**.

**Table S2** Off-target analysis of genome-edited rice mutants.

| mutant line [a] | target gene | sgRNA name | off-target site name | sequences (5′ to 3′)[b] | off-target score | location (chromosome:start) | gene locus | mutation[c] |
|---|---|---|---|---|---|---|---|---|
| *osa3′h/c5′h* & *ospmt1/2* | *OsA3′H /C5′H* | sgRNA-b | sgRNA-b-1 | G**A**ACT**A**C**G**A**G**GAGGGGCGGAAGG | 0.299 | Chr5:+3067187 | LOC_Os05g06130 | n.d. |
| | | | sgRNA-b-2 | G**C**ACT**CA**AAAGAGGGG**A**GGAAGG | 0.273 | Chr2:-25858715 | LOC_Os02g42960 | n.d. |
| *osa3′h/c5′h* | | | sgRNA-b-3 | GTAC**GC**CAA**C**GAGGGGCGGAGGG | 0.194 | Chr3:+1871092 | LOC_Os03g04080 | n.d. |

[a] Top 3 ranked potential off-target sites predicted by CRISPR-P 2.0 (Liu *et al.,* 2017) were sequenced. No potential off-target site of sgRNA-a that targets *OsFNSII* was predicted by CRISPR-P 2.0 (Liu *et al.,* 2017). [b]Bold: mismatch nucleotides compared with the sequence of sgRNA. Underlined: PAM sites. [c]n.d.: not detected.

**Table S3** Neutral sugar analysis of rice mutant cell walls.

| Sugar content (mg/g CWR) | WT | *ospmt1/2* | *osfnsII* | *osa3'h/c5'h* | *ospmt1/2 osfnsII-a* | *ospmt1/2 osfnsII-b* | *ospmt1/2 osa3'h/c5'h-a* | *ospmt1/2 osa3'h/c5'h-b* |
|---|---|---|---|---|---|---|---|---|
| Crystalline Glucose[1] | 145.3 ± 3.0[c] | **154.8 ± 5.4[b]** | **161.7 ± 2.0[ab]** | **155.2 ± 1.0[b]** | **165.6 ± 4.6[a]** | **165.3 ±0.4[a]** | **168.8 ± 4.2[a]** | **169.6 ± 1.4[a]** |
| Amorphous glucose[2] | 41.8 ± 5.8[b] | 50.1 ± 4.0[ab] | **53.4 ± 2.4[a]** | 38.4 ± 1.1[b] | 46.0 ± 4.2[ab] | 49.9 ± 3.4[ab] | 39.9 ± 1.7[b] | 43.6 ± 2.9[ab] |
| Arabinose | 39.5 ± 3.9[a] | 41.5 ± 1.6[a] | 43.1 ± 1.3[a] | 45.0 ± 1.1[a] | 44.1 ± 2.4[a] | 45.6 ± 3.5[a] | 42.8 ± 1.3[a] | 44.4 ± 2.6[a] |
| Xylose | 177.4 ± 27.0[a] | 177.8 ± 7.2[a] | 179.0 ± 9.2[a] | 171.4 ± 5.6[a] | 168.5 ± 11.8[a] | 174.9 ± 17.1[a] | 172.0 ± 5.7[a] | 172.1 ± 11.9[a] |
| Mannose | 2.4 ± 0.1[b] | 2.8 ± 0.3[ab] | 3.0 ± 0.1[ab] | 3.0 ± 0.2[ab] | 2.9 ± 0.4[ab] | **3.2 ± 0.2[a]** | 2.9 ± 0.1[ab] | **3.1 ± 0.3[a]** |
| Galactose | 19.0 ± 1.0[c] | 20.5 ± 1.2[bc] | **21.9 ± 0.6[ab]** | **23.2 ± 0.0[ab]** | **23.3 ± 1.3[a]** | **24.0 ± 1.2[a]** | **22.2 ± 0.5[ab]** | **24.3 ± 1.2[a]** |

[1]Glucose released from trifluoroacetic acid-insoluble cell wall fractions (glucose primarily from crystalline cellulose). [2]Glucose released from trifluoroacetic acid-soluble cell wall fractions (glucose primarily from amorphous cellulose and hemicelluloses). The values are means ± standard deviation from individually analyzed plants ($n$ = 3). Different letters indicate significant differences (one-way ANOVA with Tukey HSD test, $P < 0.05$). Values highlighted in bold indicate significant differences from WT. WT, wild type; *ospmt1/2*, *PMT*-deficient double-knockout line; *osfnsII*, *FNSII*-deficient single-knockout line; *osa3'h/c5'h*, *A3'H/C5'H*-deficient single-knockout line; *ospmt1/2 osfnsII-a/b*, *PMT*- and *FNSII*-deficient triple-knockout lines; *ospmt1/2 osa3'h/c5'h-a/b*, *PMT*- and *A3'H/C5'H*-deficient triple-knockout lines.

**Table S4** Peak assignment in the 2D HSQC NMR spectra of whole call wall samples.

| Labels | $\delta_C/\delta_H$ (ppm) | Assignment |
|---|---|---|
| *Lignin and hydroxycinnamate signals* | | |
| **$S_{2/6}$** | 103.9/6.74 | C2–H2 and C6–H6 in syringyl units |
| **$G_2$** | 111.0/7.03 | C2–H2 in guaiacyl units |
| **$G_{5/6}$** | 119.3/6.84, 115.8/6.83 | C5–H5 and C6–H6 in guaiacyl units |
| **$H_{2/6}$** | 128.0/7.18 | C2–H2 and C6–H6 in *p*-hydroxyphenyl units |
| **$H_{3/5}$** | 115.8/6.83 | C3–H3 and C5–H5 in *p*-hydroxyphenyl units |
| **$T_3$** | 104.9/7.04 | C3–H3 in tricin residues |
| **$T_6$** | 99.1/6.30 | C6–H6 in tricin residues |
| **$T_8$** | 94.4/6.62 | C8–H8 in tricin residues |
| **$T_{2'/6'}$** | 104.2/7.31 | C2'–H2' and C6'–H6' in tricin residues |
| **$N_{6/8}$** | 96.1/6.00, 95.2/6.04 | C2'–H2' and C6'–H6' in naringenin units |
| **$P_{2/6}$** | 130.2/7.45 | C2–H2 and C6–H6 in *p*-coumarate residues |
| **$P_{3/5}$** | 115.8/6.83 | C3–H3 and C5–H5 in *p*-coumarate residues |
| **$P_7$** | 144.9/7.47 | C7–H7 in *p*-coumarate residues |
| **$P_8$** | 113.9/6.28 | C8–H8 in *p*-coumarate residues |
| **$F_2$** | 111.1/7.34 | C2–H2 in ferulate residues |
| **$F_5$** | 115.8/6.83 | C5–H5 in ferulate residues |
| **$F_6$** | 123.3/7.11 | C6–H6 in ferulate residues |
| **$F_7$** | 145.4/7.63 | C7–H7 in ferulate residues |
| **$F_8$** | 114.2/6.52 | C8–H8 in ferulate residues |
| **$I_\alpha$** | 72.7/5.06 | Cα–Hα in β–*O*–4 units |
| **$II_\alpha$** | 87.2/5.53 | Cα–Hα in β–5 substructures |
| **OMe** | 55.7/3.69 | C–H in aromatic methoxyl groups |
| *Polysaccharide anomeric signals* | | |
| **$Gl_1$** | 103.5/4.25 | C1–H1 in (1→4)-β-D-glucopyranosyl units |
| **$X_1$** | 102.0/4.34 | C1–H1 in (1→4)-β-D-xylopyranosyl units |
| **$X'_1$** | 99.6/4.57 | C1–H1 in 2-*O*-acetyl-β-D-xylopyranosyl units |
| **$X''_1$** | 101.1/4.68 | C1–H1 in 3-*O*-acetyl-β-D-xylopyranosyl units |
| **$A_1$** | 109.4/5.22, 108.1/4.88, 107.7/5.00 | C1–H1 in α-L-arabinofuranosyl units |
| **$Ga_1$** | 105.5/4.39 | C1–H1 in (1→4)-β-D-galactopyranosyl units |
| **$U_1$** | 97.5/5.30 | C1–H1 in 4-*O*-methyl-α-D-glucuronopyranosyl units |

Measured in dimethyl sulfoxide-$d_6$/pyridine-$d_5$ (4:1, v/v). Signal assignment was based on comparison with NMR data in literature (Kim and Ralph, 2010; Mansfield *et al*., 2012; Lan *et al*., 2018; Lam *et al*., 2017; 2019; 2024; Yamamoto *et al*. 2024; Ralph et al., 2024; Ji et al., 2025).

**Table S5** Peak assignments in 2D HSQC NMR spectra of dioxane/water-soluble lignins.

| Labels | $\delta_C/\delta_H$ (ppm) | Assignment |
|---|---|---|
| $\mathbf{S_{2/6}}$ | 103.9/6.78 | C2–H2 and C6–H6 in syringyl units |
| $\mathbf{G_2}$ | 111.1/7.06 | C2–H2 in guaiacyl units |
| $\mathbf{G_{5/6}}$ | 119.4/6.87, 115.8/6.84 | C5–H5 and C6–H6 in guaiacyl units |
| $\mathbf{H_{2/6}}$ | 127.9/7.23 | C2–H2 and C6–H6 in *p*-hydroxyphenyl units |
| $\mathbf{H_{3/5}}$ | 115.8/6.84 | C3–H3 and C5–H5 in *p*-hydroxyphenyl units |
| $\mathbf{T_3}$ | 105.0/7.07 | C3–H3 in tricin units |
| $\mathbf{T_6}$ | 99.1/6.32 | C6–H6 in tricin units |
| $\mathbf{T_8}$ | 94.4/6.63 | C8–H8 in tricin r units |
| $\mathbf{T_{2'/6'}}$ | 104.3/7.36 | C2′–H2′ and C6′–H6′ in tricin r units |
| $\mathbf{N_{6/8}}$ | 96.1/6.00, 95.2/6.04 | C2′–H2′ and C6′–H6′ in naringenin units |
| $\mathbf{A_{2'/6'}}$ | 128.3/7.91 | C2′–H2′ and C6′–H6′ in apigenin units |
| $\mathbf{P_{2/6}}$ | 130.2/7.47 | C2–H2 and C6–H6 in *p*-coumarate units |
| $\mathbf{P_{3/5}}$ | 115.8/6.84 | C3–H3 and C5–H5 in *p*-coumarate units |
| $\mathbf{P_7}$ | 144.8/7.50 | C7–H7 in *p*-coumarate units |
| $\mathbf{P_8}$ | 113.9/6.32 | C8–H8 in *p*-coumarate units |
| $\mathbf{F_2}$ | 111.1/7.37 | C2–H2 in ferulate units |
| $\mathbf{F_5}$ | 115.8/6.84 | C5–H5 in ferulate units |
| $\mathbf{F_6}$ | 123.4/7.13 | C6–H6 in ferulate units |
| $\mathbf{F_7}$ | 145.0/7.65 | C7–H7 in ferulate units |
| $\mathbf{F_8}$ | 114.0/6.47 | C8–H8 in ferulate units |
| $\mathbf{I_\alpha}$ | 72.0/5.01 | Cα–Hα in β–*O*–4 units |
| $\mathbf{I_\beta}$ | 83.8/4.51, 86.2/4.21 | Cβ–Cβ in β–*O*–4 units |
| $\mathbf{I_\gamma}$ | 60.1/3.72 | Cγ–Hγ in β–*O*–4 units |
| $\mathbf{II_\alpha}$ | 87.2/5.53 | Cα–Hα in β–5 substructures |
| $\mathbf{II_\beta}$ | 53.3/3.51 | Cβ–Hβ in β–5 substructures |
| $\mathbf{II_\gamma}$ | 62.9/3.77, 63.3/3.26 | Cγ–Hγ in β–5 substructures |
| $\mathbf{III_\alpha}$ | 85.1/4.66 | Cα–Hα in resinol-type β–β substructures |
| $\mathbf{III_\beta}$ | 53.8/2.97 | Cβ–Hβ in resinol-type β–β substructures |
| $\mathbf{III_\gamma}$ | 71.0/4.13, 71.2/3.77 | Cγ–Hγ in resinol-type β–β substructures |
| $\mathbf{III'_\beta}$ | 50.3/2.67 | Cβ–Hβ in in tetrahydrofuran-type β–β substructures |
| $\mathbf{IV_\alpha}$ | 83.0/5.01 | Cα–Hα in 5–5 substructures |
| $\mathbf{IV_\beta}$ | 85.7/3.97 | Cβ–Cβ in 5–5 substructures |
| $\mathbf{V_\alpha}$ | 81.4/5.12 | Cα–Hα in β–1 substructures |
| $\mathbf{V_{\alpha'}}$ | 83.7/4.81 | Cα′–Hα′ in β–1 substructures |
| $\mathbf{V_\beta}$ | 59.9/2.84 | Cβ–Cβ in β–1 substructures |
| $\mathbf{V_{\beta'}}$ | 78.5/4.44 | Cβ′–Cβ′ in β–1 substructures |
| $\mathbf{X1_\gamma}$ | 61.7/4.16 | Cγ–Hγ in γ-free cinnamyl alcohol end-units |
| $\mathbf{X1'_\gamma}$ | 64.3/4.82 | Cγ–Hγ in γ-acylated cinnamyl alcohol end-units |
| **OMe** | 55.7/3.70 | C–H in aromatic methoxyl groups |

Measured in dimethyl sulfoxide-$d_6$/pyridine-$d_5$ (4:1, v/v). Signal assignment was based on comparison with NMR data in literature (Kim and Ralph, 2010; Mansfield *et al.*, 2012; Lan *et al.*, 2018; Lam *et al.*, 2017; 2019; 2024; Yamamoto *et al.* 2024; Ralph et al., 2024; Ji et al., 2025).
